# Myofibroblast lineage mapping and inhibiting subretinal fibrosis by targeting SMAD3 and MRTF pathways via microRNA-24 functional study

**DOI:** 10.64898/2026.03.03.709397

**Authors:** Yinga Wu, Yao Tong, Katherine G. Byrnes, Qi Zhou, Chunmin Dong, Chase Benjamin, Emma Parker, Duran Bao, Zhaoyang Ren, Chastain A. Anderson, Rafael L. Ufret-Vincenty, Yu-Guang He, Ze Zhang, David Hinkle, Jing Ma, Shusheng Wang

**Author notes:** To whom correspondence should be addressed: Shusheng Wang. The authors have declared that no conflict of interest exists.

## Abstract

Subretinal fibrosis underlies the end-stage pathogenesis of retinal diseases including age-related macular degeneration (AMD). It can disrupt retinal structure and eventually lead to legal blindness by generating contractile force, fibrotic scarring, subretinal hemorrhage, and retinal detachment. Myofibroblasts are the predominant cells critically involved in subretinal fibrosis, however, the cellular contribution to myofibroblasts remains unclear. Here we demonstrate that multiple cell lineages, including macrophages, endothelial cells (EC), retinal pigment epithelial (RPE) cells and pericytes, significantly contribute to myofibroblasts in a laser-induced subretinal fibrosis model. We found microRNA *miR-24* is significantly downregulated in the plasma of wet AMD patients. Overexpression of *miR-24* represses epithelial-mesenchymal transition (EMT), endothelial-mesenchymal transition (EndMT), and the resulting fibrosis by regulating TGF-β/SMAD3 and PAK4/LIMK2/MRTF pathways. Consistently, a combination of SMAD3 and MRTF inhibitors show superior efficacy to individual inhibitors in repressing fibrosis *in vitro* and laser-induced subretinal fibrosis *in vivo*. Together, these suggest the contribution of multiple cell-types in myofibroblast transformation in subretinal fibrosis, and repression of *miR-24*-regulated TGF-β/SMAD3 and PAK4/LIMK2/MRTF pathways in multiple cell types holds therapeutic potential for treating subretinal fibrosis in AMD and other fibrotic disorders.

## INTRODUCTION

Fibrosis is a wound healing response to mechanical- or disease-related tissue injury characterized by excess extracellular matrix (ECM) formation. Fibrotic diseases account for nearly 45% of mortality in the Western world. Several antifibrotic drugs, including Pirfenidone and Pamrevlumab, have been approved by FDA to treat fibrotic conditions, primarily in the lung and liver(1, 2). These drugs are not a cure but beneficial in slowing disease progression. Although the molecular mechanisms of fibrosis remain unclear, recent research revealed the involvement of multiple cell types, factors, and pathways. During fibrosis, inflammatory cells and other cells are recruited, activated, and proliferating in the injured regions. Myofibroblasts (activated fibroblasts) are the predominant cells that exhibit features of both fibroblasts and smooth muscle cells (SMCs) during fibrosis, including association with fibrillar collagen, formation of actin-myosin bundles (stress fibers), and exertion of contractile force(3, 4). Molecularly, transforming growth factor-β (TGF-β) is a central regulator of fibrosis, activating canonical SMAD-dependent transcriptional programs that promote mesenchymal gene expression and epithelial-mesenchymal transition (EMT)(5). Myocardin-related transcription factors (MRTFs) are required for myofibroblast differentiation by regulating myofibroblast gene programing upon actin cytoskeletal changes(6). Additional profibrotic mediators, such as connective tissue growth factor (CTGF), platelet-derived growth factor (PDGF), epidermal growth factor (EGF), and fibroblast growth factor-2 (FGF2), further amplify fibrosis through interconnected signaling networks involving MAPK, PI3K/Akt, Rho, and SMAD pathways(4, 7, 8). Diverse profibrotic stimuli, including TGF-β, CTGF, inflammatory cytokines, and mechanical cues, activate both canonical SMAD-dependent and non-canonical cytoskeletal signaling pathways. As a result, therapeutic strategies targeting a single pathway often fail due to compensatory activation of parallel cascades, highlighting the need for coordinated regulation of multiple profibrotic pathways.

Ocular fibrosis contributes to vision loss in several retinal diseases, including diabetic retinopathy, glaucoma, myopic choroidal neovascularization, and age-related macular degeneration (AMD). AMD is the leading cause of blindness in the elderly characterized by drusen deposits and retinal pigment epithelium (RPE) deterioration leading to photoreceptor loss and progressive macular degeneration. Despite the clinical success of anti-vascular endothelial growth factor (anti-VEGF) therapy in neovascular AMD (nAMD), subretinal fibrosis frequently develops and is strongly associated with poor visual outcomes(9, 10). Subretinal fibrosis arises from wound-healing responses within pre-existing neovascular membranes and disrupts retinal architecture through scar formation and tractional forces(11–13). Effective therapies targeting subretinal fibrosis remain an unmet clinical need.

Myofibroblasts are absent in healthy retinas but accumulate during subretinal fibrosis, where they deposit ECM proteins in the injured regions, including Fibronectin, Periostin, and collagens. Multiple retinal and infiltrating cell types, including Müller glia cells, RPE cells, endothelial cells (ECs), macrophages and pericytes, contribute to myofibroblasts possibly through Müller glial-mesenchymal transition (GMT), EMT, endothelial-mesenchymal transition (EndMT), macrophage-mesenchymal transition (MMT), and pericytes-myofibroblast transition (PMT), respectively(14–23). The origins of myofibroblasts, especially the relative contribution of different cell types to myofibroblasts, during subretinal fibrosis, remain unclear.

MicroRNAs (miRNAs) are endogenous small non-coding RNAs that regulate gene expression by inhibiting mRNA translation or inducing mRNA degradation(24). A list of miRNAs has been associated with fibrosis(25–28). microRNA 24 (*miR-24*) has emerged as a conserved regulator of tissue remodeling, suppressing fibrotic responses in cardiac and skeletal muscle by targeting *Furin-TGF-*β pathway and *Smad2* (27, 28). We have shown that *miR-24* represses CNV by regulating actin cytoskeletal dynamics in ECs (29). Whether *miR-24* coordinates canonical TGF-β/SMAD signaling and non-canonical cytoskeletal pathways during multicellular subretinal fibrotic reprogramming remains unknown.

Here, we quantitatively map the relative contributions of five major cell lineages to the myofibroblast population using genetic lineage tracing in a laser-induced subretinal fibrosis model. We demonstrate that subretinal fibrosis represents a multicellular reprogramming event rather than a lineage-restricted process. Mechanistically, we identify *miR-24* as an endogenous regulator that simultaneously modulates SMAD3-dependent transcriptional signaling and PAK4/LIMK2/MRTF-mediated cytoskeletal activation through distinct direct targets. Pharmacological co-inhibition of SMAD3 and MRTF phenocopies *miR-24-*mediated regulation and significantly attenuates subretinal fibrosis *in vivo*. These findings establish *miR-24* as a therapeutic lead capable of restraining fibrotic reprogramming across multiple cell lineages, highlighting the necessity of coordinated multi-pathway modulation for effective clinical intervention.

## RESULTS

### Multiple cell types contribute to myofibroblasts in the laser-injury induced subretinal fibrosis

To systematically define the cell lineages and their relative contribution to myofibroblasts during subretinal fibrosis, *in vivo* lineage-tracing studies were performed. To capture the principal retinal and infiltrating cell populations implicated in fibrotic remodeling, we selected five Cre-driver mouse lines representing RPE cells, myeloid cells, ECs, pericytes, and smooth muscle cells lineages: *VMD2^Cre^* (RPE cells), *LysM2^Cre^*(myeloid cells), *Cdh5^Cre/ERT2^* (ECs), *Pdgfr-*β*^Cre/ERT2^* (pericytes), and *Myh11^Cre/ERT2^* (SMCs). Each line was crossed with *Rosa26-tdTomato* mice for lineage-specific labeling (Fig. 1a). The non-leakiness and specificity of the Cres were validated by colocalization of tdTomato with lineage markers (Supplementary Fig.1). To assess the contribution of specific tdTomato^+^ lineage cells to the myofibroblasts, they were co-stained with cytoplasmic myofibroblast marker α-SMA at 14 days after laser injury. All evaluated lineages, including RPE cells, ECs, myeloid cells, pericytes, and SMCs, generated α-SMA^+^ myofibroblasts within the fibrotic lesions (Fig. 1b-f). Quantification using Imaris software revealed the relative contribution of different lineage cells to myofibroblasts, with 34.8% of α-SMA^+^ myofibroblasts originated from *LysM2^Cre^* (myeloid cells), 28.6% from *Cdh5^Cre/ERT2^* (EC), 18.9% from *Pdgfr-*β*^Cre/ERT2^* (pericyte), 12.0% from *VMD2^Cre^* (RPE cells), and 5.7% from *Myh11^Cre/ERT2^*(SMCs) lineages (Fig. 1g). Of note, α-SMA^+^ RPE cells were mainly localized at the edge, while α-SMA^+^ EC, pericytes, myeloid cells and SMCs were mainly localized in the middle of the lesion. When the α-SMA^+^ cell percentage in each lineage was quantified, α-SMA^+^ cells were found to constitute 10.8% of myeloid cells, 23.4% of ECs, 29.2% of Pericytes, 20.3% of RPE cells and 14.8% of SMCs, respectively (Fig. 1h). By co-immunostaining, the myofibroblast marker α-SMA was observed in ECs, pericytes, and RPE cells, but not in fibrocytes and Müller glia cells (Supplementary Fig.2). These findings suggest multiple cell types contribute to myofibroblasts during laser-induced subretinal fibrosis, with myeloid cells, ECs and pericytes as the predominant contributors. Regarding fibrotic potential, ECs, pericytes and RPE cells are more prone to myofibroblast transformation upon fibrotic stimulation.

**Fig. 1.**
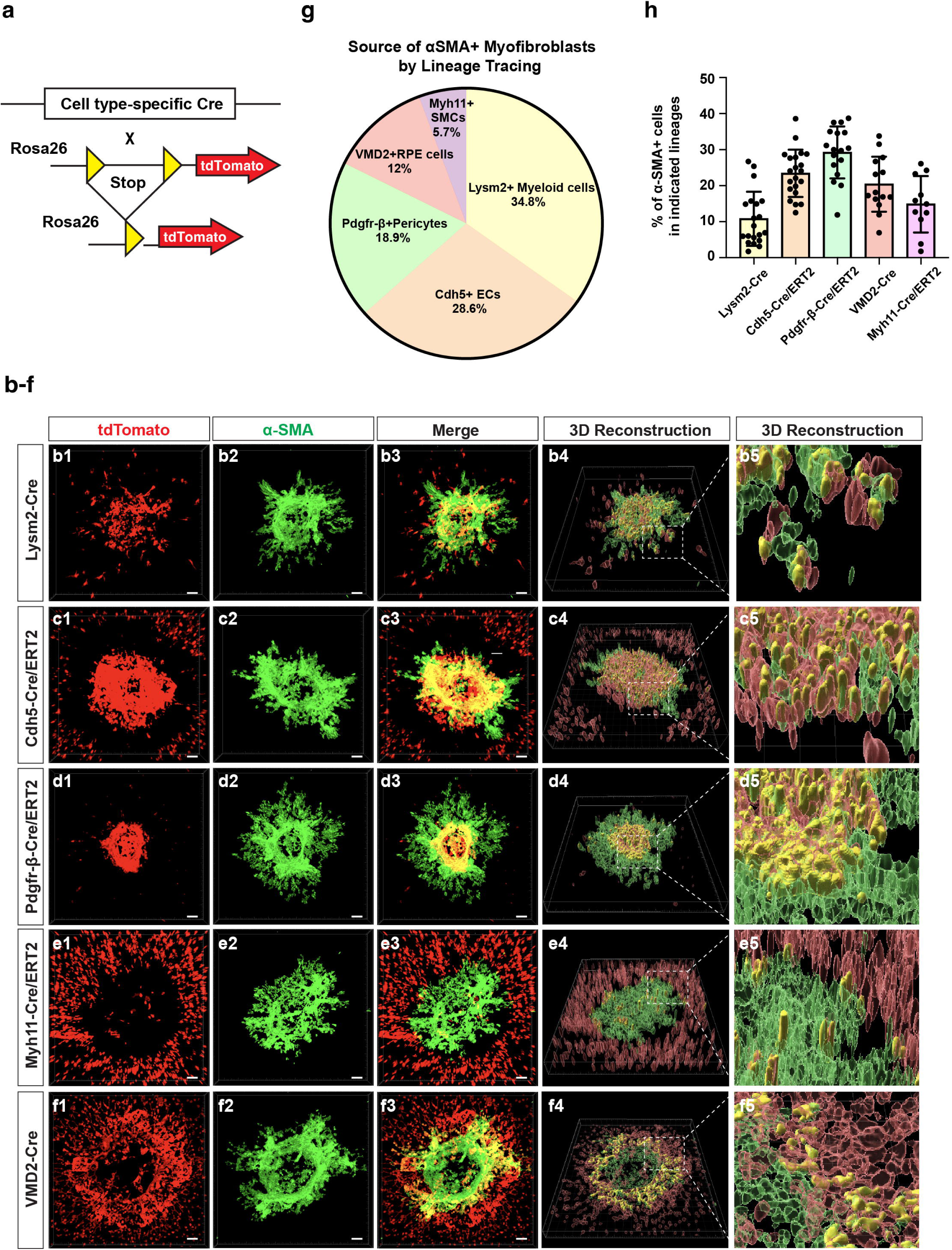
Myeloid cells, ECs, pericytes, SMCs, and RPE cells contribute to myofibroblasts in laser-induced subretinal fibrosis. **a** Schematics of specific-cell type lineage tracing using five different cell type-specific Cre mouse lines crossed with tdTomato expression in the Rosa26 locus. **b-f** Co-staining of α-SMA /tdTomato in *LysM2^Cre^;R26-tdTomato* mice **(b1-b3)** *Cdh5^Cre/ERT2^;R26-tdTomato* **(c1-c3)**, *Pdgfr-*β*^Cre/ERT2^;R26-tdTomato* **(d1-d3)**, *Myh11^Cre/ERT2^;R26-tdTomato* **(e1-e3)**, and *VMD2^Cre^;R26-tdTomato* **(f1-f3)** at 14 days after laser injury. **g** Quantification of the relative contribution of each cell-type lineage in total α-SMA^+^ myofibroblasts based on the flatmount staining in (B-F). **h** Quantification of the percentage of α-SMA^+^ cells in each cell-type lineage based on the flatmount staining in **b-f**. Each dot represents an individual lesion. Data were obtained from 21 lesions from 6 mice (*LysM2^Cre^* and *Cdh5^Cre/ERT2^*), 17 lesions from 6 mice (*Pdgfr-*β*^Cre/ERT2^*), 14 lesions from 5 mice (*VMD2^Cre^*), and 11 lesions from 5 mice (*Myh11^Cre/ERT2^*).

**Fig. 2.**
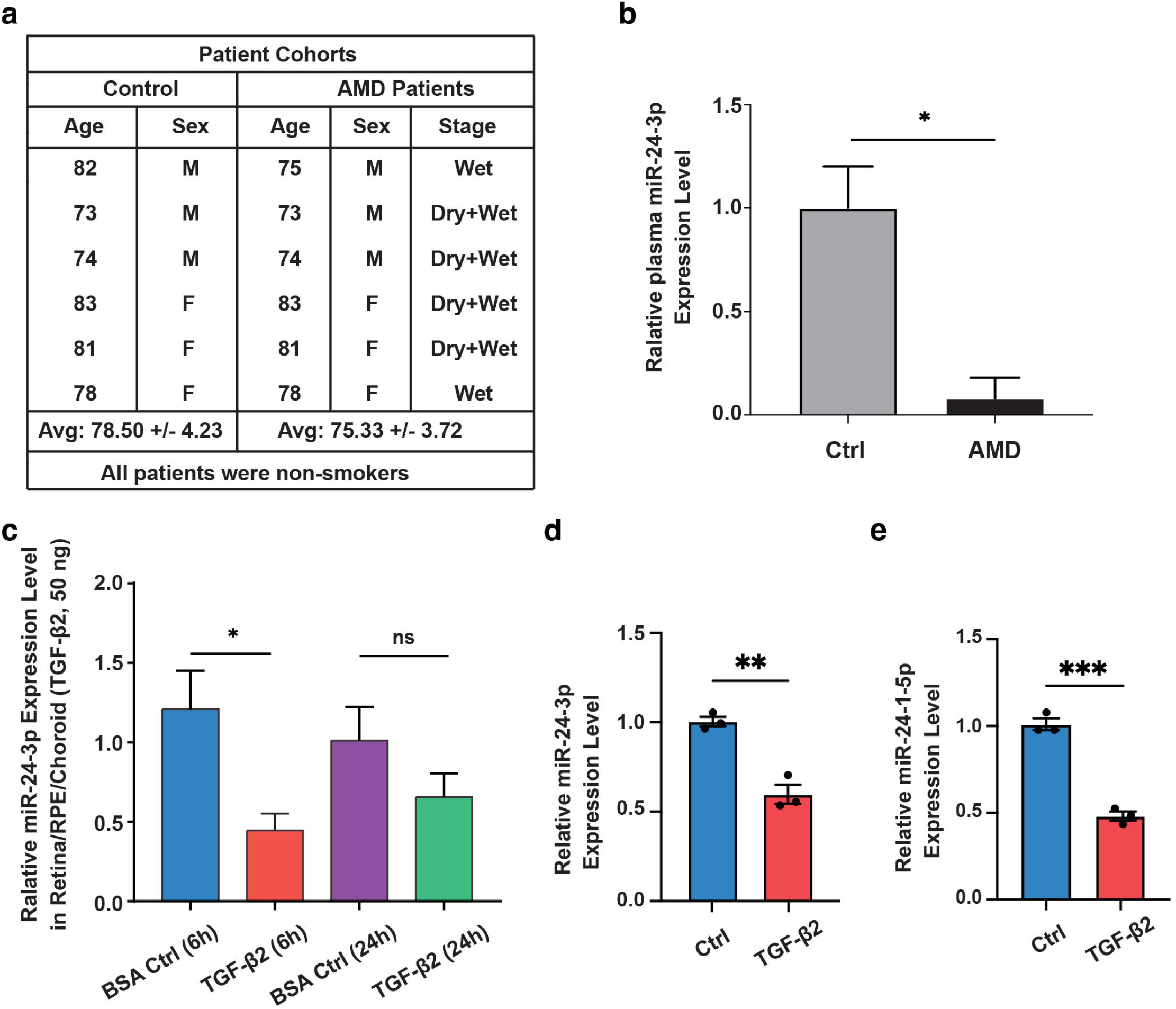
Regulation of *miR-24* expression in AMD patients, TGF-β2-treated mouse eye and ARPE-19 cells. **a** Characteristics of control and wet AMD patients. **b** Significant downregulation of *miR-24-3p* was observed in the plasma of AMD patients by miRNA profiling and confirmed by qRT-PCR. **c** Reduced *miR-24-3p* expression in Retina/RPE/Choroid at 6 and 24 hours after intravitreal injection of TGF-β2 (50ng) in mouse eye shown by RT-qPCR. **d, e** RT-PCR shows reduced expression of *miR-24-3p* (**d**) and *miR-24-1-5p* (**e**) in ARPE-19 cells after 6 hours of TGF-β2 treatment. n=3. *p < 0.05; **p < 0.01; ns, not significant.

### Regulation of *miR-24* expression by TGF-β and in the plasma of wet AMD patients

To test the involvement of *miR-24* in human nAMD, plasma microRNA profiling was performed using an Exiqon miRCURY LNA™ Universal RT microRNA PCR panel, in a group of 6 nAMD patients and 6 age-matched controls, with three males/three females in each group. The average age was 78.5 ± 4.2 for controls and 75.3 ± 3.7 for AMD patients (Fig. 2a). Patients were enrolled based on nAMD diagnosis but not specifically stratified according to the presence of subretinal fibrosis. Pooled analysis detected 398 of the 744 analyzed miRNAs in the plasma (Supplementary Fig. 3). Out of these 398 microRNAs, 52 were found to be highly up-regulated while 38 were found to be down-regulated (Supplementary Table. 1). Using individual samples, *miR-24-3p* was confirmed to be significantly downregulated in nAMD plasma (Fig. 2b).

The TGF-β pathway has been shown to suppress *miR-24* expression in skeletal muscle cells(27). To assess *miR-24* regulation in the eye, mice were intravitreally injected with TGF-β2 (50 ng). *miR-24-3p* downregulation by TGF-β2 was confirmed by qRT-PCR in retina/RPE/choroid at 6 hours, which became insignificant at 24 hours post-injection (Fig. 2c). To corroborate these findings *in vitro*, ARPE-19 cells were exposed to TGF-β2 (10 ng/mL). qRT-PCR analysis revealed that both *miR-24-3p* and *miR-24-1-5p* were significantly reduced 6 hours after treatment (Fig. 2d,e). Together, these findings identify *miR-24* as a TGF-β-responsive microRNA that is significantly reduced in patients with nAMD and in experimental models of TGF-β signaling, providing a rationale to investigate its functional role in EMT and fibrotic remodeling.

### Repression of EMT and RPE-derived fibrosis by *miR-24 in vitro*

Based on published data(27, 28) and our previous data showing the regulation of actin cytoskeletal dynamics by *miR-24*(29), we hypothesized that *miR-24* regulates EMT and subretinal fibrosis. To study the effect of *miR-24* overexpression in EMT and fibrosis, an adenovirus overexpressing *miR-24* (Ad-*miR-24*) was used to overexpress *miR-24* in ARPE-19 cells. qRT-PCR confirmed the overexpression of *miR-24*, with *miR-24-3p* increased about 3-fold and *miR-24-1-5p* increased about 18-fold relative to controls (Fig. 3a). ARPE-19 cells infected with Ad-*miR-24* or control were exposed to TGF-β2 (10 ng/mL) for 6 days to induce EMT and fibrosis. By Western blot analyses, TGF-β2 stimulation markedly increased the expression of mesenchymal marker N-Cadherin and α-SMA, whereas *miR-24* overexpression substantially attenuated this induction (Fig. 3b, c). Of note, CTGF, a downstream protein of TGF-β signaling and another key driver of EMT and fibrosis(30, 31), was almost blunted by *miR-24*, regardless of TGF-β2 treatment. These support the repression of RPE EMT by *miR-24* overexpression. To test the resultant fibrosis, ECM proteins (Fibronectin and Collagen III) were examined. Overexpression of *miR-24* repressed Fibronectin level by Western blot and the secreted Fibronectin and Collagen III by immunostaining (Fig. 3b-e). The active collagen accumulation, as visualized by collagen hybridizing peptide (F-CHP), was blunted by *miR-24* overexpression in TGF-β2-treated ARPE-19 cells (Fig. 3f, g).

**Fig. 3.**
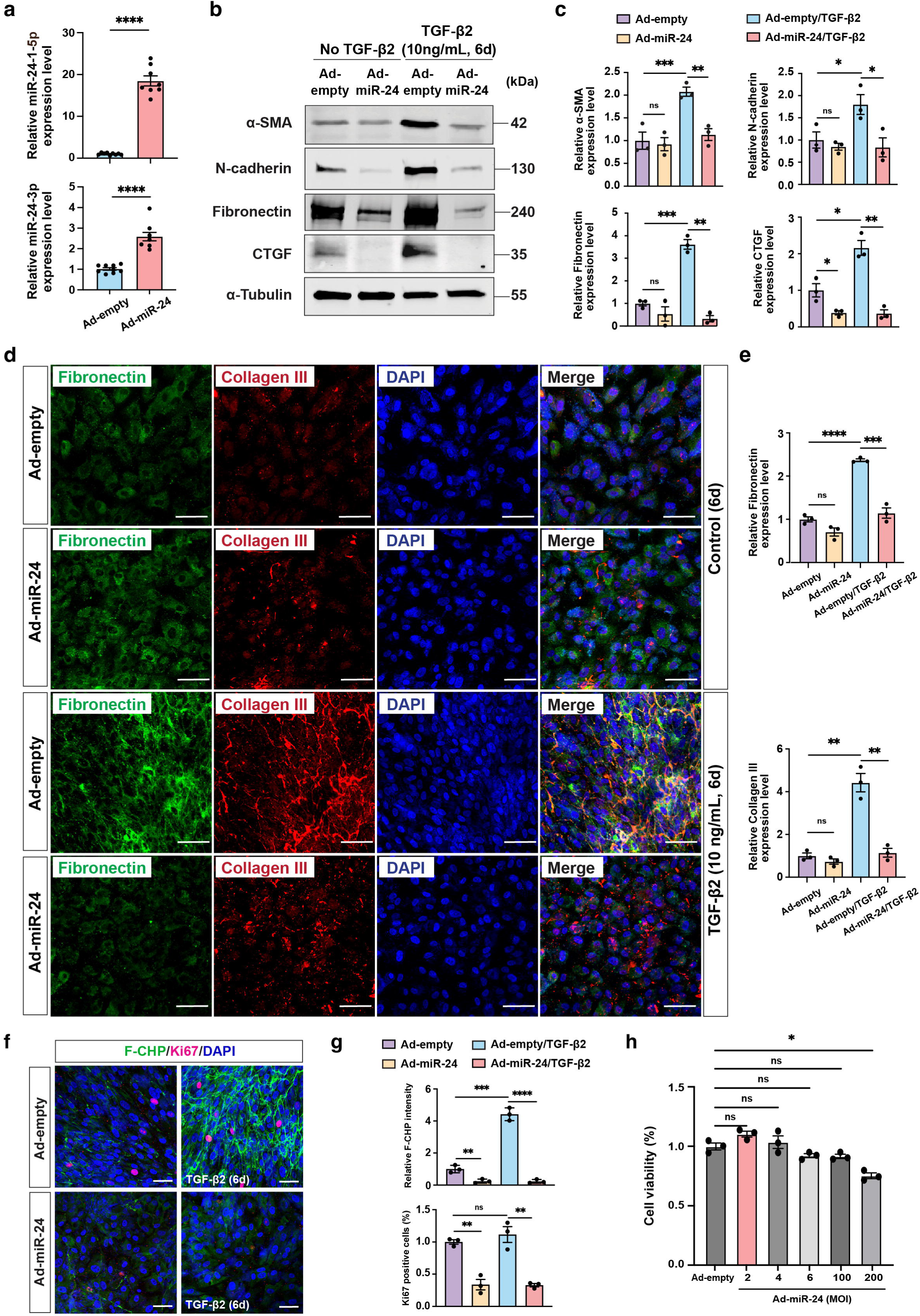
Repression of TGF-β2-induced EMT and fibrosis by *miR-24* overexpression in ARPE-19 cells *in vitro*. **a** qRT-PCR showing upregulation of *miR-24* by Adenovirus overexpressing *miR-24* (Ad-*miR-24*) in ARPE-19 cells, compared to null control (Ad-empty). MOI=2, U6 was used as an internal control. **b,c** Representative Western blots (**b**) and their quantification (**c**) showing repression of α-SMA, N-cadherin, Fibronectin, and CTGF expression by *miR-24* in ARPE-19 cells with TGF-β2 (10 ng/mL, every other day) treatment for 6 days. α-tubulin was used as a loading control. **d** Representative immunostaining indicating suppression of ECM proteins (Fibronectin and Collagen III) expression by *miR-24* in ARPE-19 cells after TGF-β2 (10ng/mL) treatment for 6 days. **e** Quantification of Fibronectin and Collagen III in (**d**). **f** Representative images showing repressed active collagen accumulation by F-CHP staining, and cell proliferation by Ki67 staining, by *miR-24* overexpression. **g** Quantification of (**f**). **h** MTT assay showing the cell viability after *miR-24* overexpression by Ad-*miR-24* at different MOIs. *p < 0.05; **p < 0.01; ***p < 0.001; ****p < 0.0001; ns, not significant.

Cell proliferation is essential for initiating EMT and driving the fibrotic cascade in myofibroblasts(32). Ki67 staining demonstrated a significant reduction in cell proliferation upon *miR-24* overexpression, regardless of TGF-β2 treatment (Fig. 3f, g), suggesting *miR-24* represses proliferation in both RPE and myofibroblast cells. Cell viability assessed by MTT assay revealed no *miR-24* cytotoxicity at the experimental dose (multiplicity of infection (MOI) of 2), although mild toxicity was observed at very high viral dose (MOI of 200), arguing against the contribution of cytotoxicity on cell proliferation (Fig. 3h).

To overcome the shortcomings of traditional ARPE-19 culture, a nicotinamide-based culture system was adopted to promote RPE differentiation, epithelial morphology and function(33). ARPE-19 cells were cultured in MEM-Nam medium for two weeks prior to infection with Ad-*miR-24*. Differentiated ARPE-19 cells developed mature cell-cell junctions shown by ZO-1 immunostaining (Supplementary Fig. 4a). To induce EMT and fibrotic responses, differentiated ARPE-19 cells were treated with TGF-β2 (10 ng/mL), TNF-α (10 ng/mL), or their combination (referred to as TNT) for 6 days. Successful induction of EMT and fibrosis by TNT was indicated by the elongated RPE shape shown by ZO-1 staining, and immunostaining using Collagen III antibody and F-CHP staining (Fig. 4a, b, Supplementary Fig. 4). TGF-β2 or TNF-α alone was less efficient (Supplementary Fig. 4). *miR-24* overexpression attenuated TNT-induced EMT and fibrosis, as evidenced by the blunted F-CHP signal and Collagen III staining, and lack of RPE elongation (Fig. 4a, b, Supplementary Fig. 4b-d). Collectively, these results demonstrated that *miR-24* exerts anti-fibrotic effect by suppressing TGF-β2 and TNT-induced EMT, ECM protein production and cell proliferation.

**Fig. 4.**
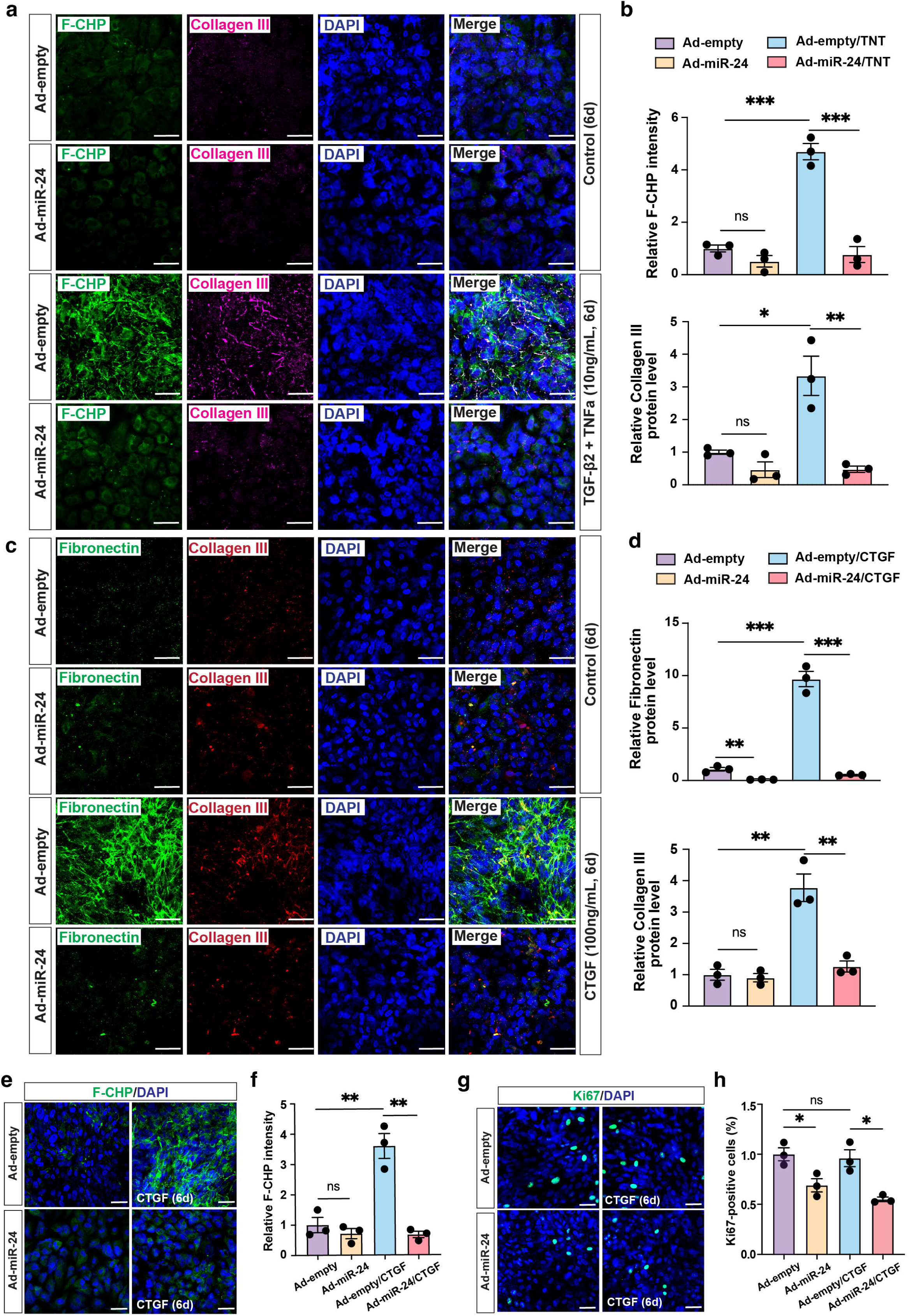
*miR-24* overexpression represses TGF-β2/TNF-α-induced EMT in differentiated ARPE-19 cells and CTGF-induced EMT in ARPE-19 cells. **a,b** Representative immunostaining showing (**a**) and quantification (**b**) showing repression of TGF-β2/TNF-α (TNT)-induced fibrosis (F-CHP and Collagen III staining) by *miR-24* overexpression in ARPE-19 cells cultured with MEM-Nam medium for 2 weeks. **c, d** Representative images (**c**)and quantification (**d**) indicating repression of Fibronectin and Collagen III expression by *miR-24* overexpression in ARPE-19 cells treated with CTGF (100 ng/mL, every other day) for 6 days. **e, g** Representative images showing blunted F-CHP staining (**e**), and repressed cell proliferation by Ki67 staining (**g**), by *miR-24* overexpression in ARPE-19 cells treated with CTGF (100 ng/mL) for 6 days. **f, h** Quantification of (**e**) and (**g**). n=3. *p < 0.05; **p < 0.01; ***p < 0.001, ****p < 0.0001; ns, not significant.

### Repression of CTGF-induced fibrosis *in vitro* and TGF-β/TNF-α-induced subretinal fibrosis by *miR-24 ex vivo*

CTGF is another strong inducer of EMT and fibrosis, functioning by amplifying pro-fibrotic signaling via upregulating factors including TGF-β (34, 35). It has been implicated in nAMD, with elevated expression observed in CNV lesions and advanced fibrotic regions(36). To determine whether *miR-24* can repress CTGF-induced EMT and fibrosis in ARPE-19 cells, ARPE-19 cells were infected with Ad-*miR-24* for three days, followed by stimulation with CTGF (100 ng/mL) every other day for 6 days. CTGF markedly induced EMT and fibrosis, as shown by the robust upregulation of Fibronectin and Collagen III by immunostaining (Fig. 4c, d). *miR-24* overexpression blunted this response, which was also confirmed by F-CHP staining (Fig. 4e, f). In addition, *miR-24* overexpression reduced CTGF-induced cell proliferation as shown by Ki67 staining (Fig. 4g, h). These support *miR-24* as a potent suppressor of CTGF-induced fibrosis.

To validate our findings in a physiologically relevant model, we modified an *ex vivo* explant system to culture laser-injured RPE/choroid(37). RPE/choroid explants from laser-induced injury were cultured on PVDF membranes and then infected with Ad-empty or Ad-*miR-24* for 2 days, followed by TNT treatment for 4 days. *miR-24* overexpression in the RPE/Choroid explants was confirmed by RT-qPCR (Fig. 5a). *Ex vivo* culture of TNT-treated RPE/choroid laser spots generated strong EMT, and fibrotic response shown by flatmount α-SMA and Fibronectin staining (Fig. 5b, c). *miR-24* overexpression markedly reduced TNT-induced EMT and fibrotic changes, suggesting an anti-fibrotic role in an *ex vivo* model of subretinal fibrosis.

**Fig. 5.**
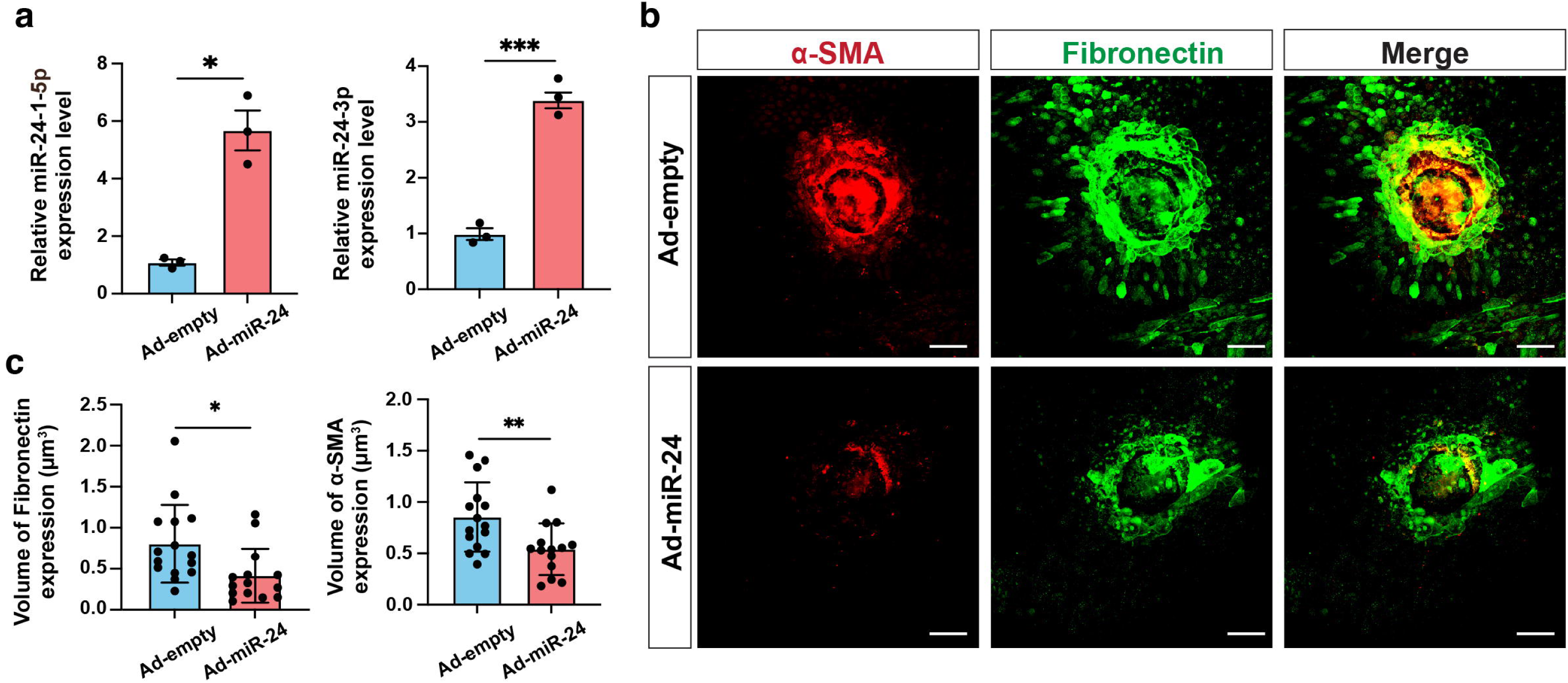
Suppression of α-SMA and Fibronectin by *miR-24* in laser-induced subretinal fibrosis *ex vivo*. **a** RT-qPCR showing overexpression of *miR-24* in RPE/Choroid explants treated with Ad-*miR-24* for 3 days. **b** Flatmount staining showing reduced expression of α-SMA and Fibronectin by *miR-24* overexpression in a laser-induced subretinal fibrosis *ex vivo* model. (**c**) Quantification in (C). n=3. *p < 0.05; **p < 0.01.

### Repression of EndMT and EC-derived fibrosis by *miR-24 in vitro*

EndMT has been implicated in tissue fibrosis. Consistent with a recent study, we showed that ECs transdifferentiate into myofibroblasts after laser injury (Fig.1) (16). To investigate the role of *miR-24* in regulating EndMT and fibrosis, HUVECs and MS1 cells were infected with Ad-*miR-24* for 3 days and treated with a cytokine cocktail containing TGF-β2 (5 ng/mL), IL-1β (100 ng/mL), and TNF-α (5 ng/mL) every other day for 6 days. Western blot analysis confirmed *miR-24* overexpression suppressed mesenchymal and fibrotic protein expression, including N-cadherin and fibronectin (Fig. 6a, b). Immunofluorescence staining further confirmed that *miR-24* overexpression decreased the secretion of ECM proteins, including Fibronectin and Collagen III (Fig. 6c, d). This was also confirmed in TNT-treated MS1 cells by F-CHP staining (Supplementary Fig. 5a, b). Together, these results support a repressive role for *miR-24* in EndMT and resultant fibrosis.

**Fig. 6.**
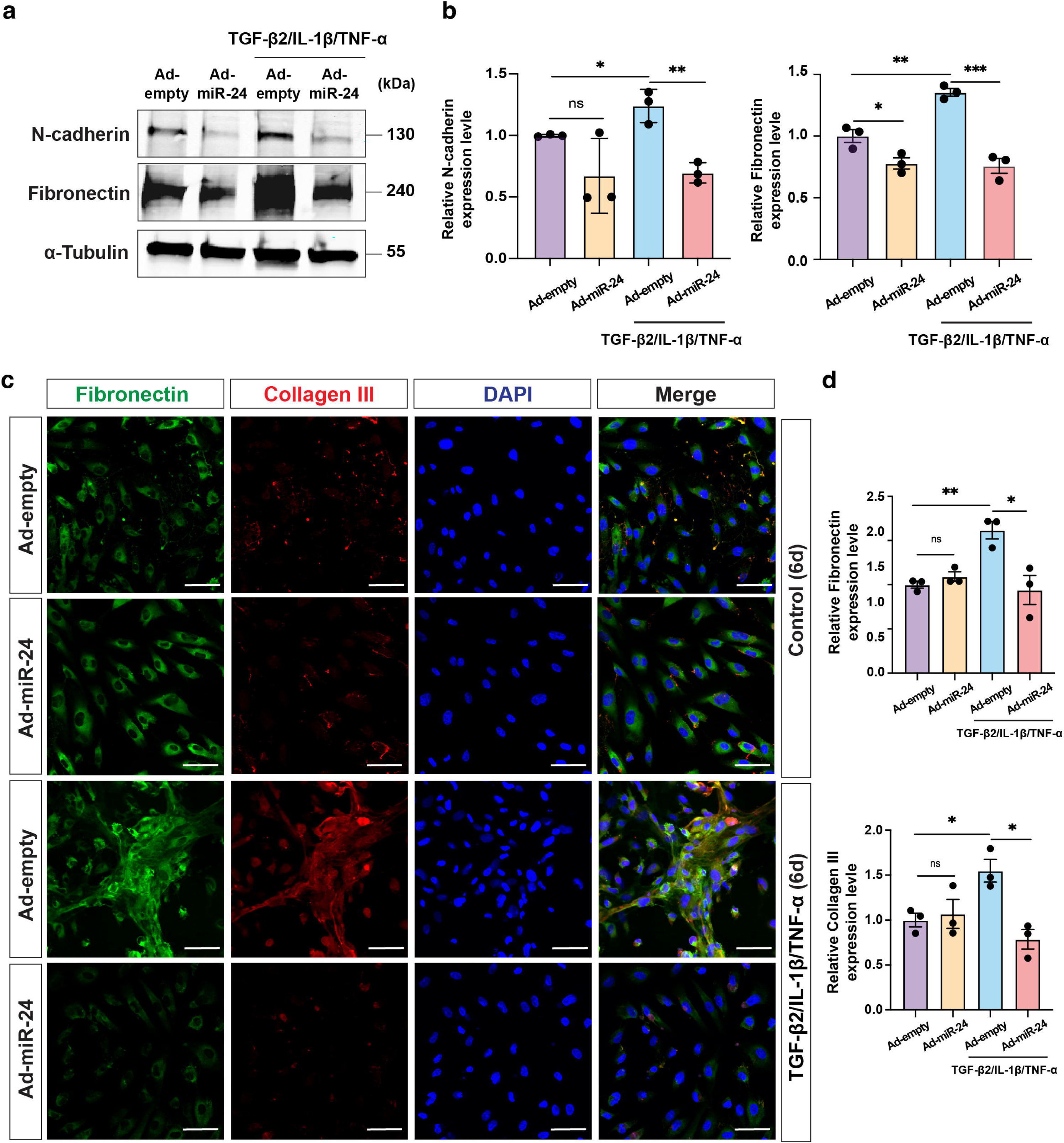
Suppression of EndMT and fibrosis by *miR-24* overexpression in TGF-β2/TNF-α/IL-1β-induced EndMT and fibrosis in HUVECs. **a** Western blots showing inhibition of N-cadherin and Fibronectin expression by Ad-*miR-24* in HUVECs treated with TGF-β2/TNF-α/IL-1β for 6 days. α-Tubulin served as loading controls. **b** Quantification of (**a**). **c** Immunostaining showing blunted Fibronectin and Collagen III expression by *miR-24* overexpression in similar samples to (**a**)**. d** Quantification of (**c**). n=3. *p < 0.05; **p < 0.01; ***p < 0.001; ****p < 0.0001; ns, not significant.

### Target genes and pathways regulated by *miR-24* in EMT and fibrosis

To investigate the global transcriptional effects of *miR-24*, RNA sequencing (RNA-seq) was performed in differentiated ARPE-19 cells under the following conditions: Ad-empty control or Ad-*miR-24* with or without profibrotic stimulation (TNT). *miR-24* overexpression elicited highly consistent transcriptional effects, with 1475 and 1493 genes upregulated [Fold of Change (FC) >2] and 767 and 659 genes downregulated (FC>2) under basal and TNT conditions, respectively. Strong concordance in gene regulation was observed across contexts (Pearson *r* = 0.84, *P* < 0.0001; Supplementary Fig. 6a). Over-representation analysis (ORA) of differentially expressed genes revealed that *miR-24* induces a broad reprogramming of the RPE transcriptome. Among the top 10 downregulated pathways, *miR-24* overexpression suppressed mitotic cell cycle, DNA replication, and cell proliferation, together with major fibrotic signaling axes including TGF-β and PI3K-Akt signaling, indicating coordinated inhibition of stress-induced proliferative and profibrotic programs. In contrast, the top 10 upregulated pathways were enriched for cytoprotective and stress-adaptive responses, including defense response, response to wounding, and response to oxidative stress, as well as negative regulation of mesenchymal cell differentiation, consistent with a robust anti-EMT transcriptional signature (Fig. 7a).

**Fig. 7.**
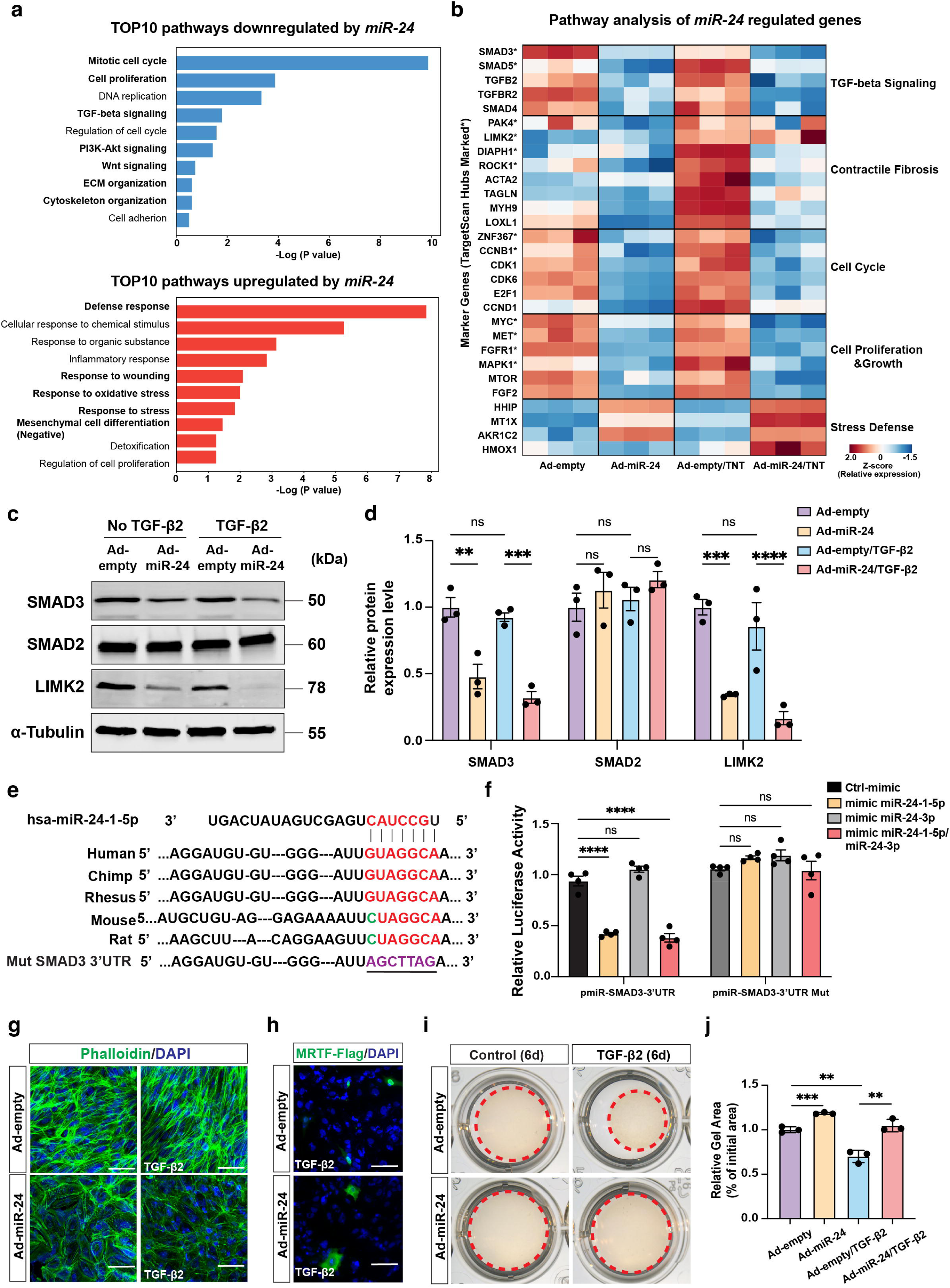
*miR-24* Overexpression downregulates the expression of EMT markers by regulating TGF-β/SMAD3 and LIMK2/MRTF pathways *in vitro*. **a** Pathway enrichment analysis of genes downregulated (blue) or upregulated (red) by *miR-24* overexpression in ARPE-19 cells, showing the top 10 pathways ranked by -log_10_(*P* value). **b** Heatmap of relative expression (Z-score) of selected marker genes and predicted or validated *miR-24* targets upon Ad-*miR-24* infection, and with or without TNT treatment. Genes are grouped by functional categories. Asterisks (*) indicate TargetScan-identified *miR-24* target hub genes. **c** Repression of SMAD3, LIMK2, but not SMAD2 protein expression by *miR-24* overexpression in ARPE-19 cells, with or without TGF-β2 treatment (10 ng/mL) for 6 days. **d** Quantification in (**c**)**. e** Predicted binding sites of *miR-24* with target genes in human, chimpanzee, rhesus monkey, mouse, and rat. The sequences in the 3’UTR of SMAD3 complementary to the miRNA seed regions are shown in red. The mutant sequence of the 3’UTR of SMAD3 is shown in blue. **f** *miR-24-1-5p* but not 3p mimic repressed SMAD3 3’-UTR luciferase activity, which was blunted in *miR-24* binding mutant. **g** Repression of stress fiber formation by *miR-24* overexpression in APRE-19 cells, with or without TGF-β2 treatment for 6 days, visualized with Phalloidin staining. **h** Blocking of FLAG-MRTFA nuclear translocation by *miR-24* in MRTF-transfected cells treated with TGF-β2 (10 ng/mL) for 24hr, visualized by FLAG staining. **i, j** Representative images (**i**) and quantification (**j**) showing reduced gel contractility in *miR-24*-overexpressing ARPE-19 cells with or without TGF-β2 treatment (10 ng/mL) for 4 days. Red dashed circles indicate the gel area. Data are presented as relative gel area (% of initial area). n=3. **p < 0.01; ***p < 0.001; ****p < 0.0001; ns, not significant.

To further delineate the relationship between *miR-24*-regulated transcriptional networks and *miR-24* target genes, *miR-24-1-5p* and *3p* target genes were downloaded from Targetscan (version 7.0), and cross compared with the differentially expressed gene by *miR-24* overexpression. 20-27% of the *miR-24-1-5p* and *miR-24-3p* target genes were downregulated by *miR-24* overexpression, suggesting both *miR-24-1-5p* and *miR-24-3p* are actively hitting their targets (Supplementary Table 2). We curated predicted and experimentally validated *miR-24* targets together with representative downstream effector genes involved in fibrosis, contractility, cell-cycle control, and stress response (Fig. 7b and Supplementary Fig. 6b). Consistent with pathway enrichment results, *miR-24* overexpression coordinately repressed profibrotic and proliferative networks under both basal and TNT conditions, including TGF-β signaling components (*SMAD3*, *SMAD5*, *TGFBR2*), contractile fibrosis regulators (*LIMK2*, *PAK4*, *DIAPH1*, *ROCK1*, *ACTA2*, *TAGLN*, *MYH9*, *LOXL1*), and key cell-cycle drivers (*CCNB1*, *CDK1*, *CDK6*, *E2F1*, *MYC*, *FGFR1*, *MAPK*1, *MTOR*), consistent with the observed phenotypes by us and others(38). This suppression extended beyond predicted targets to downstream pathway effectors, indicating network-level regulation rather than isolated gene-specific effects. In contrast, cytoprotective genes (*HMOX1*, *AKR1C2*, MT1X) were preserved or induced in *miR-24*-treated cells, particularly under stress. Together, these findings show that *miR-24* reshapes the RPE transcriptome by targeting central regulatory nodes and attenuating profibrotic and proliferative programs. Genome-wide association studies have identified more than 34 independent loci conferring AMD risk, including *CFI*, *ABCA1*, *VEGFA*, and *TGFBR1*, which are dysregulated in AMD patient tissues (Supplemental Table 3)(39). Consistent with patient data, TNT treatment did not significantly alter *ABCA1* expression but downregulated *CFI* while upregulating *VEGFA* and *TGFBR1* in Ad-empty cells (Supplementary Fig. 6c). *miR-24* overexpression reversed these disease-associated expression patterns under TNT conditions, restoring homeostatic genes and suppressing angiogenic and profibrotic signaling.

To further investigate *miR-24* functional mechanisms in fibrosis, potential EMT- and fibrosis-related *miR-24* target genes and pathways were examined. *LIMK2* is a proven *miR-24-3p* target gene in ECs(29). The *SMAD2* and *SMAD3* genes were potential *miR-24* targets by HITS-CLIP sequencing(40, 41), with *SMAD2* confirmed as a *miR-24* target in skeletal muscle(27). Western blot analysis confirmed the repression of *LIMK2* and *SMAD3*, but not *SMAD2* protein expression by *miR-24* overexpression in ARPE-19 cells, regardless of TGF-β2 treatment (Fig. 7c, d). Similar results were also observed in HUVEC, suggesting *SMAD3*, but not *SMAD2*, is a *miR-24* target in RPE and ECs (Supplementary Fig. 7). A 6-7 nucleotide sequence complementary to the *miR-24-1-5p* seed region was observed in the *SMAD3* 3’-UTRs, which is conserved in multiple species including humans, chimpanzees, rhesus monkeys, mice, and rats (Fig. 7e). To validate *SMAD3* as a direct *miR-24-1-5p* target, a human *SMAD3* 3’-UTR fragments containing this target site was cloned downstream of a luciferase reporter and tested for luciferase activity after *miR-24* mimic transfection in 293T cells. Co-transfection with *miR-24-1-5p* mimics, but not *miR-24-3p*, significantly reduced luciferase activity (Fig. 7f). This effect was abolished when the *miR-24-1-5p* targeting site was mutated, confirming *SMAD3* as a direct target of *miR-24*.

To investigate the impact of *miR-24* overexpression on TGF-β/LIMK2 signaling, actin cytoskeletal dynamics and its downstream signaling were examined. TGF-β promotes actin cytoskeletal remodeling by activating LIMK2, therefore increasing stress fiber formation and nuclear translocation of MRTF by reducing G-actin levels(42). *miR-24* overexpression in ARPE-19 cells inhibited TGF-β2-induced stress fiber formation, as shown by Phalloidin staining (Fig. 7g, Supplementary Fig. 8a), decreased F-actin/G-actin ratio (Supplementary Fig. 8b), and suppressed MRTF-A nuclear translocation (Fig. 7h). Actin cytoskeletal dynamics are critical for cellular contractility, a hallmark of differentiated myofibroblast in fibrosis(43). TGF-β2-induced cell contractility was almost blunted by *miR-24* overexpression in ARPE-19 cells, as demonstrated by gel contraction assay (Fig. 7i, j). Together, these data indicate *miR-24* inhibits fibrosis by targeting SMAD3 and LIMK2 in the TGF-β/SMAD3 and PAK4/LIMK2/MRTF pathways.

### Dual inhibition of SMAD3 and MRTF represses fibrosis *in vitro* and *in vivo*

To further elucidate *miR-24*’s functional mechanism, pharmacological inhibitors of *miR-24* targeted pathways were used to determine whether they could mimic the anti-fibrotic effects of *miR-24*. SIS3 (Sigma, S0447) is a specific inhibitor of SMAD3 functioning by blocking SMAD3 phosphorylation and its interaction with SMAD4. CCG-203971 (MedChemExpress, HY-108361) is a MTRF inhibitor functioning by blocking MRTF nuclear accumulation. ARPE-19 cells were primed with SIS3 at 10 μM, and/or CCG-203971 at 20 μM, prior to TGF-β2 (10 ng/mL) treatment for 6 days to induce fibrosis. The combination of SIS3 and CCG-203971 treatment resulted in almost complete inhibition of the fibrotic phenotypes, shown by blunted Fibronectin secretion by immunostaining and F-CHP staining, mimicking the effect of *miR-24* overexpression (Fig. 8a, b). Individual inhibitors also partially repressed fibrotic phenotypes, with better effects at higher dosages and CCG-203971 being more potent than SIS3 (Supplementary Fig. 8c-e).

**Fig. 8.**
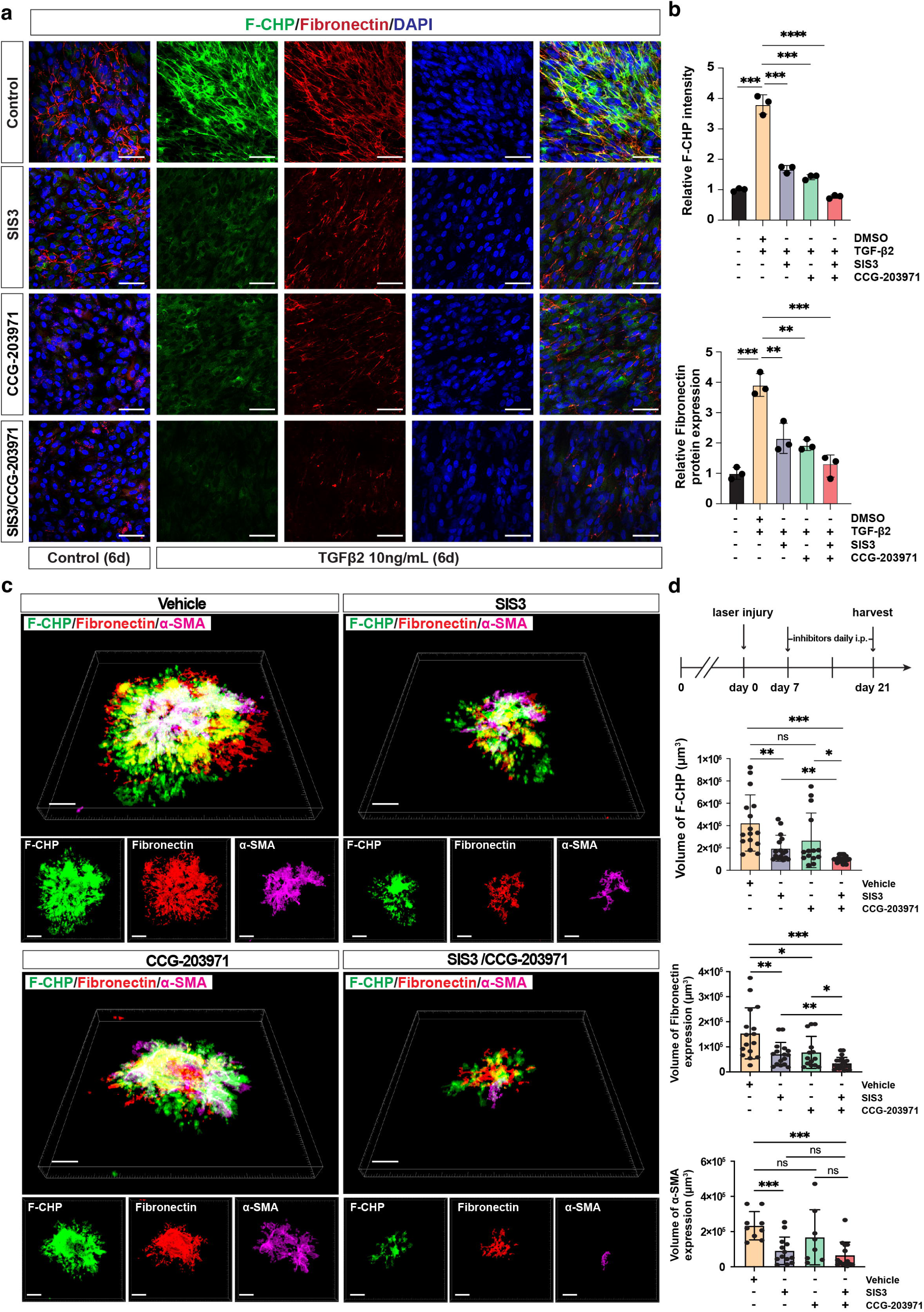
Inhibition of SMAD3 and MRTF activity represses laser-induced subretinal fibrosis *in vitro* and *in vivo*. **a** Representative immunostaining images showing repressed expression of Fibronectin and F-CHP staining by SMAD3 inhibitor (SIS3) at 10 μM or MRTF-A inhibitor (CCG-203971) at 20 μM, individually or in combination, in TGF-β2-treated ARPE-19 cells. **b** Quantification of (**a**). n=3. **c** Experimental timeline for evaluation of dual inhibition of SMAD3 and MRTF activity in the laser-induced subretinal fibrosis model. Representative three-dimensional reconstructions images showing RPE/choroid flatmount staining for F-CHP (green), Fibronectin (red), and α-SMA (magenta) are shown, with individual channels displayed below. **d** Quantification of (**c**). n = 6 mice per group, with 3 laser spots per eye. Eyes with hemorrhage after laser photocoagulation were excluded from analysis. *p < 0.05; **\***\*p < 0.01; ***p < 0.001; ****p < 0.0001; ns, not significant.

To evaluate the therapeutic efficacy of SMAD3 and MRTF inhibition in laser-induced subretinal fibrosis, C57BL/6J mice were subjected to laser injury and treated with SIS3 and/or CCG-203971, or vehicle control. Inhibitors were administered daily by intraperitoneal injection from 7-21 days post injury, when CNV is peaked and subretinal fibrotic response is active (Fig. 8c, d). Dual inhibition of SMAD3 and MRTF markedly repressed laser-induced EMT and subretinal fibrosis as shown by RPE/choroid flatmount staining with α-SMA, fibronectin, and F-CHP (Fig. 8c, d). Quantitative volumetric analysis of fluorescence signal revealed a 77% reduction in accumulated active collagen (F-CHP□ volume; vehicle vs SIS3/CCG-203971: mean = 423,875μm^3^ vs 100,316μm^3^), an 88% decrease in α-SMA expression (α-SMA^+^volume; vehicle vs SIS3/CCG-203971: mean = 233,111μm^3^ vs 65,531μm^3^), and a 79% reduction in Fibronectin deposition (Fibronectin□ volume; vehicle vs SIS3/CCG-203971: mean = 153,925μm^3^ vs 34,493μm^3^) compared with vehicle-treated controls. In contrast, SIS3 or CCG-203971 alone resulted in less efficient repression of subretinal fibrosis. These indicate that concurrent blockade of the two major *miR-24*-regulated pathways, TGF-β/SMAD3 and LIMK2/MRTF, is required for robust suppression of myofibroblast activation and extracellular matrix accumulation during subretinal fibrosis. To determine whether earlier intervention could similarly limit fibrotic remodeling, we administered SIS3/CCG-203971 immediately following laser injury and continued daily until day 14 (Supplementary Fig. 9a-c). Consistently, early dual inhibition significantly reduced fibrotic burden, shown by a 60% decrease in F-CHP-positive collagen, a 79% reduction in α-SMA expression, and a 73% reduction in fibronectin deposition compared to controls (Supplementary Fig. 9a-c), indicating early intervention has similar effect and therefore is not required for inhibiting laser-induced subretinal fibrosis. In addition, combined inhibition of SMAD3 and MRTF significantly suppressed CNV formation, reducing IB4-positive lesion volume by approximately 87% compared with controls (Supplementary Fig. 9c, d). Collectively, these findings demonstrate that inhibition of the *miR-24*-regulated TGF-β/SMAD3 and LIMK2/MRTF pathways effectively suppresses EMT and fibrotic responses *in vitro* and attenuates both CNV formation and subretinal fibrosis *in vivo*.

## Discussion

Subretinal fibrosis underlies the pathology in multiple retinal diseases, including AMD. It develops in nAMD patients regardless of anti-VEGF treatment and is associated with CNV recurrence and poor visual outcomes. The molecular mechanisms and cellular contribution of subretinal fibrosis remain unclear. By systematic lineage tracing and functional study of *miR-24*, our study provides new insight into subretinal fibrosis. Our major findings include: (1) multiple cell-types, including ECs, myeloid cells, pericytes, RPE cells, and SMCs, contribute to myofibroblasts in subretinal fibrosis; (2) *miR-24* expression is downregulated in the plasma of nAMD patients and by TGF-β2 in the retina and RPE cells; (3) Overexpression of *miR-24* represses TGF-β2 or CTGF-induced EMT in RPE cells and TGF-β2/IL-1β/TNF-α induced EndMT in ECs, as well as the resultant fibrosis in *in vitro* and *ex vivo* models; (4) *miR-24* promotes an anti-fibrotic and anti-proliferation transcriptional program and functions by repressing TGF-β/SMAD3 and PAK4/LIMK2/MRTF signaling by targeting LIMK2 (*miR-24-3p* target) and SMAD3 (*miR-24-1-5p* target); (5) A combination of SMAD3 and MRTF inhibitors repress fibrotic response *in vitro* and laser-induced CNV and subretinal fibrosis *in vivo*. Collectively, these findings support a model in which subretinal fibrosis is driven by myofibroblasts from multiple lineages and sustained by the convergence of canonical TGF-β/SMAD3 and non-canonical PAK4/LIMK2/MRTF signaling (Supplementary Fig. 10). Within this framework, *miR-24* emerges as an endogenous brake that restrains myofibroblast transformation from multiple lineages.

The cellular origins of myofibroblasts in subretinal fibrosis remain incompletely understood. Here, through systematic and stringent genetic lineage tracing, we demonstrate that ECs, myeloid cells, and pericytes, represent major sources of α-SMA^+^ myofibroblasts following laser injury, whereas RPE cells and SMCs contribute to a lesser extent. (Fig. 1). Previous studies have implicated macrophages as a potential source of myofibroblasts based primarily on marker co-staining; co-localization of F4/80 and α-SMA has been observed in both experimental models and human nAMD samples, suggesting macrophage involvement. A lineage-tracing study in the laser CNV model further demonstrated that myeloid-lineage cells do not transdifferentiate into endothelial cells; however, their potential contribution to α-SMA^+^ myofibroblasts was not evaluated(44). Likewise, RPE contribution to myofibroblasts was supported by marker co-staining (α-SMA and cytokeratin) in surgically excised AMD-associated CNV membranes(45), whereas an early-stage lineage-tracing study (day 5 CNV) did not detect substantial RPE-derived myofibroblasts(44). Pericyte involvement has been proposed using Col1α1-GFP transgenic mice, in which TGF-β2 enhanced GFP□ cell accumulation and apparent transition toward a myofibroblast phenotype in laser-induced CNV(46). However, because Col1α1 is expressed in fibroblasts and myofibroblasts in addition to pericytes, Col1α1-GFP reporters cannot unequivocally trace pericyte fate during fibrosis. In contrast, tamoxifen-inducible *Pdgfr-*β*^Cre/ERT2^* permits pre-injury, lineage-specific labeling of pericytes, allowing more definitive quantification of their contribution to myofibroblasts. Consistent with a recent report(16), we observe a substantial endothelial contribution and extend these findings by providing a systematic comparison of the relative contribution of major retinal and infiltrating lineages. Collectively, our results support a model in which subretinal fibrosis reflects a coordinated multicellular reprogramming event rather than a lineage-restricted phenomenon. This conceptual shift broadens the therapeutic landscape beyond RPE-EMT, underscoring the need to address vascular and inflammatory cell plasticity, particularly EndMT, PMT, and MMT in subretinal fibrosis. These findings suggest that effective antifibrotic therapies for nAMD will likely require targeting multiple cellular sources of myofibroblasts rather than focusing exclusively on RPE-derived

EMT. Our study did not fully assess several additional cell types that have been proposed to participate in retinal fibrotic remodeling. Müller glia have been proposed as a potential source of myofibroblasts in subretinal fibrosis, as cultured Müller cells can lose glutamine synthetase (GS) and GFAP expression while acquiring α-SMA expression(47). Although technical limitations prevented lineage tracing in *Pdgf*α*^Cre^;R26-tdTomato* mice due to impaired fundus visualization following laser injury (Supplementary Fig. 11a, b), marker-based analyses don’t support a substantial Müller glial contribution. At day 21 after laser injury, we observed minimal co-localization of GS and α-SMA (Supplementary Fig. 2). Furthermore, co-staining for RPE65 and GS revealed low-level GS expression in the RPE layer under baseline conditions, suggesting that the rare GS□/α-SMA□ cells likely represent RPE-derived myofibroblasts rather than Müller glia migrating from the neural retina. These findings suggest that Müller glia cells contribute minimally, if at all, to myofibroblast population in this model. Circulating fibrocytes have been detected in CNV membranes and fibrotic lesions, representing another putative source(48, 49). However, triple immunostaining for Collagen I, CD45, and α-SMA revealed little to no Collagen I□/CD45□/α-SMA□ cells at day 21 following laser injury, suggesting minimal fibrocyte-to-myofibroblast transition in our model (Supplementary Fig. 2). However, the lack of fibrocyte-specific genetic tools limits definitive lineage tracing, and future studies employing adoptive transfer or bone marrow-based approaches will be required to clarify their contribution. Resident choroidal fibroblasts may represent another potential source. Evaluation of their involvement would require fibroblast-specific inducible systems, such as *Tcf21*^Cre/ERT^. Additionally, analysis across multiple time points may further refine understanding of stage-dependent lineage contributions.

One limitation of the present study is that genetic lineage tracing was performed using separate Cre driver lines, preventing simultaneous tracking of multiple cellular origins within the same lesion. Consequently, lineage contributions were estimated from independent experiments and should be interpreted as relative rather than definitive contribution of the specified lineages to the total α-SMA□ myofibroblast population. If the involvement of additional cell populations such as choroidal fibroblasts is confirmed, the relative ratio could change. Despite this limitation, the overall α-SMA□ myofibroblast burden was sufficiently comparable across experiments to provide meaningful insight into the relative importance of different cellular sources. Furthermore, our conclusions were supported by immunofluorescence co-staining analyses and an independent myofibroblast lineage-tracing approach using Postn-MCM mice. Consistent with previous reports identifying Periostin as a marker of activated myofibroblasts, the vast majority of Periostin-lineage-labeled cells co-expressed α-SMA□ within fibrotic lesions (Supplementary Fig. 12), providing independent validation that α-SMA□ cells represent the major myofibroblast population in this model. While single-cell RNA sequencing and spatial transcriptomics can reveal cellular states and spatial relationships, they cannot directly establish lineage origin and therefore complement rather than replace genetic lineage tracing. Interestingly, RPE-derived α-SMA cells were preferentially localized at the lesion border, whereas myofibroblasts derived from other lineages were more prominent within the lesion core, suggesting the existence of spatially and potentially functionally distinct myofibroblast subpopulations during subretinal fibrosis.

An increasing list of miRNAs have been shown to regulate fibrotic diseases, including *miR-21*, *miR-24*, *miR-29*, *miR-200* family members, primarily in the lung, heart, kidney and liver(26–28, 50–55). For example, *miR-21* promotes fibrosis by enhancing TGF-β signaling, whereas genetic deletion or anti-*miR-21* treatment reduces fibrosis in multiple models(25, 26, 50, 51). *miR-29* is markedly downregulated in renal, pulmonary, and cardiac fibrosis, leading to increased extracellular matrix expression, and restoration of *miR-29* using chemically modified mimics attenuates fibrosis in the lung and skin(53, 54, 56–58). The *miR-200* family, particularly *miR-200a* and *miR-200c*, is downregulated in pulmonary fibrosis in mouse models and human samples, and delivery of *miR-200* mimics or precursors reduces pulmonary, renal, and peritoneal fibrosis *in vivo*(55, 59). *miR-24* is a multi-functional miRNA shown to regulate apoptosis, angiogenesis and fibrosis. Lentiviral or mimic-mediated overexpression of *miR-24* suppresses fibrosis in the heart and skeletal muscle by targeting Furin and SMAD2, respectively(27, 28). However, miRNA functions and targets could be tissue- and context-dependent (e.g., *miR-24* does not target SMAD2 in RPE or HUVECs in our study), underscoring the need for validation across multiple models before clinical translation.

Here, we found that *miR-24* is reduced in the plasma of AMD patients and is suppressed by TGF-β2 in the retina and cultured RPE cells (Fig. 2). Because the human cohort was not specifically characterized for subretinal fibrosis, the reduction in circulating *miR-24* should be interpreted as an AMD-associated finding. However, our mechanistic studies in multiple fibrosis models support a direct antifibrotic role for *miR-24*. Nevertheless, our mechanistic studies in multiple *in vitro*, *ex vivo*, and *in vivo* fibrosis models support a direct role for *miR-24* in regulating profibrotic responses. Our previous study that *miR-24* inhibits CNV by targeting PAK4, LIMK2, and DIAPH1 prompted us to explore its function in subretinal fibrosis(29). Given that multiple cell types contribute to myofibroblast formation, current work is focused on EMT and EndMT as key transitional processes. Although undifferentiated ARPE-19 cells exhibit certain mesenchymal-like characteristics, TGF-β2 treatment further induced robust expression of mesenchymal and fibrotic markers, consistent with previous reports. Importantly, *miR-24* not only suppressed TGF-β2-induced EMT but also reduced basal expression of several mesenchymal genes in ARPE-19 cells, supporting a direct role for *miR-24* in maintaining epithelial homeostasis. *miR-24* overexpression suppressed EMT in ARPE-19 cells induced by TGF-β2 or CTGF, and EndMT in ECs induced by TGF-β2, TNF-α, and IL-1β. These effects were accompanied by reduced expression of mesenchymal markers, diminished extracellular matrix deposition, and decreased active collagen accumulation (Fig. 3, 4, 6). To address the limitations of the conventional ARPE-19 model, we further validated these findings in a long-term differentiated ARPE-19 culture system with preserved tight-junction integrity and in an *ex vivo* RPE/choroid explant model derived from laser-injured eyes, both of which more closely recapitulate physiological conditions (Fig. 4, 5, Supplementary Fig. 4). Although direct *in vivo* delivery of *miR-24* mimics in the laser-induced fibrosis model remains technically challenging in our hand, pharmacological inhibition of two major downstream pathways regulated by *miR-24* significantly attenuated CNV and subretinal fibrosis *in vivo*. Our observation that *miR-24* suppresses both TGF-β2-induced CTGF expression and CTGF-driven fibrosis is of particular interest, as CTGF blockade has been shown to reduce subretinal fibrosis in laser injury models and a CTGF-targeting RNAi therapy is currently under clinical evaluation (NCT02599064)(60, 61). The ability of *miR-24* to inhibit fibrotic phenotypes induced by diverse stimuli across multiple cell types positions it as a promising therapeutic candidate for subretinal fibrosis in nAMD. Because *miR-24* suppresses both CNV and fibrosis, it may offer a unique therapeutic advantage by simultaneously targeting the neovascular and fibrotic components of disease. Given that subretinal fibrosis frequently develops despite successful anti-VEGF treatment, a single agent capable of suppressing both processes could provide broader and more durable therapeutic benefit than approaches directed at either pathology alone. Future studies will determine whether *miR-24* similarly regulates fibrosis from additional myofibroblast-producing lineages, including macrophages and pericytes.

Transcriptomic analyses further revealed that *miR-24* exerts broad, network-level control over profibrotic and proliferative programs in RPE cells. RNA-seq revealed highly consistent *miR-24*-mediated transcriptional responses under both basal and profibrotic stress conditions, indicating robust regulatory activity. Pathway analyses showed coordinated suppression of cell-cycle progression and major fibrotic signaling pathways, including TGF-β and PI3K-Akt signaling, alongside activation of cytoprotective and stress-adaptive programs. Importantly, *miR-24* reversed the expression patterns of several independent AMD risk genes that are dysregulated in patient tissues linking its transcriptional effects to clinically relevant disease pathways. Together, these data support a model in which *miR-24* reprograms stressed RPE and ECs away from a proliferative, mesenchymal-like state toward a more homeostatic and stress-resilient phenotype.

Mechanistically, *miR-24* regulates fibrosis by targeting key nodes that control actin cytoskeletal remodeling through TGF-β/SMAD3 signaling and PAK4/LIMK2/MRTF pathway. SMAD3, a central mediator of TGF-β signaling, was identified as a novel direct target of *miR-24-1-5p* in human cells (Fig. 7). SMAD3 is a well-established driver of EMT and fibrosis, and *Smad3*-deficient mice exhibit broad resistance to fibrotic injury in multiple organs(62, 63). In a proliferative vitreoretinopathy model, *Smad3^-/-^* RPE cells maintain a monolayer morphology without EMT marker upregulation(64), underscoring the importance of this pathway in retinal fibrosis. In parallel, we have previously shown that *miR-24* targets LIMK2 in ECs, a key effector in the Rho/PAK4/LIMK2/MRTF-A axis(29). Here, we show that *miR-24* also represses LIMK2 in RPE cells (Fig. 7c, d), resulting reduced actin polymerization, diminished cellular contractility, and impaired MRTF nuclear translocation (Supplementary Fig. 8, Fig. 7g, h). As LIMK2 promotes stress fiber assembly and MRTF-A functions as a mechanosensitive SRF coactivator that drives myofibroblast gene expression(65), these findings place *miR-24* upstream of two major profibrotic signaling axes. Consistent with this axis, genetic or pharmacological inhibition of MRTF reduces fibrosis in multiple tissues, including ocular contexts(6, 66, 67), local delivery of MRTF inhibitors such as CCG-222740 or CCG-203971 reduced postoperative scarring in a clinically relevant rabbit model of glaucoma filtration surgery, and intravitreal administration of CCG-1423 attenuated photocoagulation-induced Collagen I expression *in vivo*(68). Importantly, we found that combined inhibition of SMAD3 and MRTF suppressed subretinal fibrosis more potently *in vitro* and *in vivo* (Fig. 8c, d, Supplementary Fig. 9). Our data confirmed the role of *miR-24* in ocular fibrosis, extending the previous work on *miR-24* in cardiac and skeletal muscle fibrosis. These findings reveal strand-specific functions of *miR-24*, with *miR-24*-3p targeting LIMK2 and *miR-24-1-5p* targeting SMAD3, thereby enabling coordinated regulation of parallel profibrotic pathways. *SMAD2* was not regulated by *miR-24* in human RPE and EC cells, despite prior reports in murine systems(27), highlights species- and context-specific differences in miRNA-target interactions.

Subretinal fibrosis arises from complex interactions among inflammatory, angiogenic, and mechanical cues. Although many antifibrotic approaches focus on individual cytokines such as TGF-β or CTGF, fibrosis can also be driven by extracellular matrix remodeling, mechanical stress, and actin cytoskeletal signaling. Epithelial injury is required to initiate EMT, as TGF-β alone is insufficient to activate myofibroblast programs in intact epithelia(69). In a two-hit model, disruption of epithelial contacts first activates Rho/Rac signaling and drives MRTF nuclear translocation (first hit), while subsequent TGF-β signaling provides a second hit that enables full transcriptional activation of myofibroblast genes(70). SMAD3 interacts with MRTF and cooperatively regulates the timing of MRTF-mediated transcription during EMT(70, 71), highlighting functional cooperation between these pathways.

Consistent with this model, analysis of LINCS L1000 CRISPR knockout consensus signatures identified 81 genes co-regulated by SMAD3 and MRTF-A, as well as 417 genes uniquely associated with either pathway(72)(Supplementary Fig. 13a). KEGG enrichment analysis revealed complementary but distinct signaling signatures: shared targets were enriched in cytoskeletal organization (Supplementary Fig. 13b), SMAD3-specific genes were associated with canonical TGF-β, AGE-RAGE, and ECM-related pathways, whereas MRTF-A-specific genes were enriched in Ras, complement/coagulation, and vascular remodeling pathways (Supplementary Fig. 13c, d, Supplementary Table 4). Genes downregulated by *miR-24* significantly overlapped with both SMAD3- and MRTF-A-regulated transcriptional signatures, indicating that *miR-24* suppresses both shared and pathway-specific profibrotic programs (Supplementary Fig. 13a). These findings support a model in which *miR-24* coordinately attenuates cytokine-driven (SMAD3-dependent) and mechanically reinforced (MRTF-dependent) fibrotic signaling, providing mechanistic support for its broad antifibrotic activity. Although direct *in vivo* delivery of *miR-24* was not evaluated, the consistency of the antifibrotic effects observed across *in vitro*, *ex vivo*, transcriptomic, and downstream pathway inhibition studies supports the biological relevance of *miR-24* in regulating subretinal fibrosis.

Therapeutic restoration of *miR-24*, either through viral gene delivery or chemically modified miRNA mimics, may represent a promising strategy for treating subretinal fibrosis in nAMD. By coordinately suppressing SMAD3- and MRTF-dependent profibrotic pathways across multiple cell types, *miR-24* has the potential to exert broader antifibrotic effects than approaches targeting a single downstream pathway. Alternatively, combined SMAD3/MRTF inhibition may provide a pharmacological strategy to recapitulate key antifibrotic effects of *miR-24*. Future studies are needed to directly compare the dual inhibitors and miR-24 overexpression.

Our previous study demonstrated that *miR-24* suppresses laser-induced CNV *in vivo*(29), whereas the present study establishes its antifibrotic activity across multiple cell types and models. Because subretinal fibrosis frequently develops despite successful VEGF suppression, a single agent capable of targeting both pathological processes could provide broader and more durable therapeutic benefit than approaches directed at either process alone. Given that anti-VEGF agents and SMAD3/MRTF-targeted therapies primarily address distinct pathological components of disease, combinatorial regimens integrating these modalities may further improve outcomes, particularly in patients at high risk for macular scarring. While MRTF inhibition alone has shown limited efficacy in suppressing CNV(73), combined inhibition of SMAD3 and MRTF significantly reduces CNV formation (Supplementary Fig. 9). As MMT and PMT play a prominent role in myofibroblast transformation (Fig. 1), future studies are needed to test the effect of *miR-24* in MMT and PMT.

An important limitation of the present study is that the in vivo therapeutic experiments were performed using pharmacological inhibition of SMAD3 and MRTF rather than direct delivery of *miR-24*. Therefore, although our data support the conclusion that *miR-24* exerts antifibrotic effects through these pathways, they do not establish that *miR-24* replacement itself is sufficient to suppress subretinal fibrosis *in vivo*. Future studies utilizing optimized viral vectors, chemically modified miRNA mimics, or nanoparticle-based delivery systems will be required to directly evaluate the therapeutic efficacy of *miR-24* in animal models of subretinal fibrosis.

In sum, our findings identify *miR-24* as a central regulator of subretinal fibrosis that acts by coordinately suppressing TGF-β/SMAD3 and LIMK2/MRTF signaling. As targeting a single signaling pathway frequently fails to halt fibrosis due to redundancy and compensatory activation of alternative profibrotic circuits, *miR-24*-based therapy represents a “one drug/multiple targets” paradigm that can broadly suppress fibrosis by simultaneously modulating key pathways, potentially overcoming limitations of current single-target approaches, including incomplete efficacy, dose-limiting toxicity, and drug resistance.

## MATERIALS AND METHODS

### Human samples and miRNA profiling

Blood samples from wet AMD patients or age-matched controls were collected in the clinics in the Department of Ophthalmology at UTSW. All subjects were examined by slit-lamp biomicroscope, optical coherence tomography, fundus photography, and fluorescein angiography. Patient information was de-identified and passed to researchers with patient information limited to age, sex, smoking status, and stage of AMD. 6 nAMD patients and 6 control subjects (3 males and 3 females per group) were included. All AMD subjects had wet AMD; 4 also had dry AMD. Patients were not specifically selected or stratified based on the presence of subretinal fibrosis, and therefore fibrosis status was not available for analysis. Smokers were excluded to minimize confounding effects.

Blood samples were collected in 10 mL heparin tubes and centrifuged at 500×g for 10 min at 4 °C. Plasma was carefully collected above the buffy coat, transferred to RNase-free tubes, and centrifuged again at 500×g for 10 min at 4 °C. Supernatants were aliquoted and either processed immediately or stored at −80 °C. RNA isolation from plasma was carried out using TRIzol™ (Invitrogen) according to the protocol of the manufacturer with the following modifications: isopropanol was added at a 1:1 ratio with the initial volume of TRIzol™ and the isopropanol precipitation was carried out at -80 °C overnight as described(29). For the Exiqon miRCURY LNA™ Universal RT microRNA PCR panels, RNA extracts from the six patients in each group were pooled prior to cDNA synthesis to produce two samples. cDNA was synthesized with the miRCURY LNA™ Universal RT microRNA PCR Kit (Exiqon) per the manufacturer’s protocol. The Exiqon system consists of two 384 well qPCR plates containing primers for 744 microRNAs of interest, along with inter-plate calibrators and housekeeping controls. qPCR was performed on an ABI 7900HT system. Data were analyzed using GenEx software, and following inter-plate calibration, Ct values were normalized to the mean Ct of all expressed microRNAs. Validation of selected miRNAs in individual plasma samples were performed by RT-qPCR for miRNA using miRCURY LNA SYBR Green PCR kit (Qiagen, Cat Log. 339345).

### Animals and treatments

Mouse lines are as follows: *Postn^MCM/+^*, B6.129S-Postntm2.1(cre/Esr1*)Jmol/J, Jax stock No: 029645; *Myh11^Cre/ERT2^*, B6.FVB-Tg(Myh11-icre/ERT2)1Soff/J, Jax stock No: 019079; *Pdgfr-*β*^Cre/ERT2^*, B6.Cg-Tg(Pdgfrb-cre/ERT2)6096Rha/J, Jax stock No: 029684; *Cdh5^Cre/ERT2^*mouse line was provided by Dr. Stryder Meadows from Tulane university; *Rosa26-tdTomato* mice line was provided by Dr. Maria Galazo from Tulane university; *LysM2^Cre^*mouse line was provided by Dr. Xuebin Qin from Tulane university; *VMD2^Cre^* mouse line was provided by Dr. Alfred S. Lewin from University of Florida. Mice (8-12-week-old) were used in the studies on laser-induced subretinal fibrosis. Both sexes were used in the experiments.

Tamoxifen was used to induce Cre recombinase activity, including MerCreMer (MCM) protein or CreERT2 protein. For *Myh11^Cre/ERT2^* mice, *Cdh5^Cre/ERT2^* mice, and *Pdgfr-*β*^Cre/ERT2^* mice, tamoxifen (Sigma, T5648), dissolved in a 90% corn oil/10% Ethanol solution by shaking overnight at 37°C, was administered to mice at 75 mg tamoxifen/kg body weight for 5 consecutive days by intraperitoneal (IP) injection at least 7 days before laser injury to induce subretinal fibrosis. For *Postn^MCM^* mice, tamoxifen (20 mg/mL) were given by IP injection every other day after laser injury. To induce Cre expression in *VMD2^Cre^* mice, doxycycline-containing chow (Inotiv, TD00502; 200 mg/kg) was administered to nursing dams from postnatal day 1 to day 14, leading to Cre activation in the pups.

### Laser-induced subretinal fibrosis model

Laser-induced subretinal fibrosis was performed using an established laser-induced CNV model(29). Briefly, 6-8-week-old male C57BL/6J mice were anesthetized via intraperitoneal injection of ketamine (100□mg/kg) and xylazine (5□mg/kg). Pupils were dilated with 1% tropicamide (Alcon Laboratories, Fort Worth, TX), and a topical analgesic (0.5% proparacaine) was applied. Gonak (2.5% hypromellose solution; Akorn, Lake Forest, IL) was used to maintain corneal hydration. Fundus imaging was conducted using a Micron III retinal imaging system (Phoenix Research Laboratories, Pleasanton, CA), and laser photocoagulation was performed using a 532□nm YAG laser (Meridian Merilas, Thun, Switzerland) coupled to the Micron III via a laser injector. Parameters were optimized to 1000 mW (≈150 mW output), 100 ms duration, and 100 µm spot size. Three burns were applied per eye. Bruch’s membrane rupture was confirmed by bubble formation. Following laser photocoagulation, the lubricant eye gel was gently removed and replaced with erythromycin ophthalmic ointment to prevent infection. Samples were collected at 14 and 21 days after laser injury for fibrotic marker staining. For lineage-tracing experiments, 5-8 mice were included in each Cre-driver group. Laser photocoagulation was performed with three burns per eye. Eyes and individual lesions were evaluated according to predefined quality-control criteria adapted from established recommendations for laser-induced CNV studies(74). Exclusion criteria included unsuccessful laser burns without Bruch’s membrane rupture, excessive hemorrhage, fused or morphologically abnormal lesions, lesion distortion caused by animal movement during laser application, tissue damage during dissection or processing, and lesions classified as outliers based on predefined morphological criteria. Following quality-control assessment, all eligible lesions were quantified. Depending on lesion exclusion, a total of 11–21 lesions were analyzed per experimental group.

To evaluate the efficacy of SMAD3 and MRTF-A inhibitors in laser-induced subretinal fibrosis, two experimental models were employed. In the first model, C57BL/6J mice were subjected to laser injury at 12 weeks of age. Beginning 7 days after laser photocoagulation, mice received daily intraperitoneal injections of the SMAD3 inhibitor SIS3 (Sigma, S0447; 2.5 mg/kg), the MRTF-A inhibitor CCG-203971 (MedChemExpress, HY-108361; 100 mg/kg), or both compounds for an additional 14 days. For drug preparation, SIS3 and CCG-203971 were initially dissolved in DMSO at 50 mg/mL and 250 mg/mL, respectively. Stock solutions were diluted to working concentrations using stepwise solvent systems, with PEG-300, Tween-80, and sterile saline added sequentially and mixed thoroughly after each addition. The final formulation for SIS3 consisted of 1% DMSO, 40% PEG-300, 5% Tween-80, and 54% sterile saline (5 μL/g body weight; e.g., 100 μL for a 20 g mouse), whereas CCG-203971 was formulated in 5% DMSO, 40% PEG-300, 5% Tween-80, and 50% sterile saline (8 μL/g body weight; e.g., 160 μL for a 20 g mouse). For combination treatment, the two compounds were prepared separately and administered as two sequential intraperitoneal injections. Corresponding vehicle formulations without inhibitors were used as controls. Animals were randomly assigned to four treatment groups: (1) vehicle (DMSO/saline), (2) SIS3, (3) CCG-203971, and (4) SIS3/CCG-203971.

In the second model, 12-week-old C57BL/6J mice were subjected to laser photocoagulation and received daily intraperitoneal injections of SIS3 (2.5 mg/kg) and CCG-203971 (100 mg/kg), prepared separately and administered as two sequential injections, beginning immediately after laser injury and continuing daily through day 14. Drug preparation and vehicle controls were identical to those described above. Animals were randomly assigned to two treatment groups: (1) vehicle (DMSO/saline) and (2) SIS3/CCG-203971.

### Plasmid construction and reporter assays

The human SMAD3 3’-untranslated region (3’-UTR) was amplified from a GeneCopoeia template plasmid (Smad3-3UTR-MT05b) by polymerase chain reaction and cloned into the pmiR-REPORT™ luciferase vector (Ambion). To assess the specificity of *miR-24* targeting, a mutant SMAD3 3’-UTR reporter was generated by site-directed mutagenesis at the predicted *miR-24* binding site using overlap extension PCR. Briefly, two overlapping fragments were amplified using primer pairs Smad3-3’UTR-F/Smad3-3-UTR-Mut-R and Smad3-3’UTR-Mut-F/Smad3-3’UTR-R, followed by a second PCR to generate the full-length mutant 3’-UTR. Then, the mutant fragment was subcloned into the pmiR-REPORT™ vector. All constructs were verified by restriction enzyme digestion and Sanger sequencing. Primers used are: Smad3-3’UTR-F: ttaattaatcgGAGCTCAGACCTGAAGGCTACTTCTAGGA; Smad3-3’UTR-R: ttaattaatcgAAGCTTAAATGCCATTCATACAACAGG; Smad3-3’UTR-Mut-F: GATGTGTGGGATTagcttagAACCCACCT; Smad3-3’UTR-Mut-R: AGGTGGGTTctaagctAATCCCACACATC. Luciferase reporter assays were performed as described(29). In brief, 293T cells were transfected with the pmiR-REPORT™ luciferase vector containing the wild-type or mutant SMAD3 3′-UTR, together with *mir*Vana^®^ *miR-24-3p* mimic (Thermo Fisher Scientific, Assay ID MC10737), *mir*Vana^®^ *miR-24*-1*-5p* mimic (Thermo Fisher Scientific, Assay ID MC12902), or their combination. A β-galactosidase plasmid was used as an internal control. Luciferase activity was measured using the Luciferase Assay System (Promega, Cat. #E1500) according to the manufacturer’s instructions.

### Adenovirus and infection

Adenovirus generation and transduction were performed as described(29). pShuttle-*miR-24*vector was generated by cloning human Pre-*miR-24* (using primers: 5’-CGCGGTGAACTCTCTCTTGT-3’) and 5’-CCCAGCTCTCCTGAGCCT-3’) sequences from race-ready human heart cDNA into the pShuttle-CMV vector. Then, this vector was recombinantly inserted into an adenoviral vector and generated Adenovirus overexpressing *miR-24* (Ad-*miR-24*) using the AdEasy system(75). Empty adenovirus (Ad-empty) was used as a control in the experiments. The titer of Ad-*miR-24* or Ad-empty was measured using Adeno-X qPCR titration kit (Takara, 632252). Briefly, viral DNA was extracted from the preparation and amplified using quantitative PCR (qPCR) with specific primers provided in the kit. The resulting Ct values were compared to a standard curve to calculate viral genome copy number, allowing for accurate and sensitive quantification of the viral titer.

### Cell culture and treatment

ARPE-19 cells, HUVEC cells, and MS1 cells were used in the experiments. ARPE-19 cells (ATCC® CRL-2302™) were cultured in Dulbecco’s modified Eagle’s medium with nutrient mixture F-12 (DMEM/F-12) with phenol red (HyClone, SH3028401), supplemented with 10% fetal bovine (HyClone), and 1% Penicillin-Streptomycin (10,000 U/mL)(Gibco, 15140122). Human Umbilical Vein Endothelial Cells (HUVEC) (Lonza, C2517A) were cultured in Endothelial Cell Growth Medium-2 (EGM-2) (Lonza, CC3162). MILE SVEN 1 (MS1) cells (ATCC® CRL-2279™) were cultured in Cytiva HyClone™ Dulbecco’s High Glucose Modified Eagles Medium (HyClone, SH30022LS), supplemented with 10% fetal bovine (HyClone), and 1% Penicillin-Streptomycin (10,000 U/mL) (Gibco, 15140122). AD-293 cells, used for adenovirus production, were cultured in Cytiva HyClone™ Dulbecco’s High Glucose Modified Eagles Medium (HyClone, SH30022LS), supplemented with 10% fetal bovine (HyClone), 1% Penicillin-Streptomycin (10,000 U/mL) (Gibco, 15140122). For long-term RPE culture model that can differentiate human RPE cells using nicotinamide(33), ARPE-19 cells were cultured in MEM alpha medium (Gibco, 12571063) supplemented with GlutaMAX™ Supplement (ThermoFisher, 35050061), 1% fetal bovine (HyClone), 1% Penicillin-Streptomycin (10,000 U/mL)(Gibco, 15140122), 1% N1 supplement (sigma, N6530), MEM Non-Essential Amino Acids Solution (ThermoFisher, 11140050), taurine (0.25 mg/mL) (T8691; sigma), hydrocortisone (20 ng/mL) (Sigma, H6909), triiodo-thyronin (0.013 ng/mL) (Sigma, T6397), and nicotinamide (10mM) (Sigma, N0636). Cells were maintained at 37°C in a humidified atmosphere containing 5% CO□, with the medium replaced every 2 days for 14 days. Cells at passages 2 to 6 were used for all *in vitro* experiments.

ARPE-19 cells were seeded in 12-well or 6-well plates at approximately 80% confluency. Three days following transduction with Ad-*miR-24* at MOI of 2, cells were treated with TGF-β2 (10 ng/mL) or CTGF (100 ng/mL) every other day for a total duration of 6 days to induce EMT. For long-term differentiated ARPE-19 culture model, EMT and fibrosis were induced by treatment with TGF-β2 (10 ng/mL), TNF-α (10 ng/mL), or their combination, administered every other day for 6 days. Similarly, HUVECs and MS1 cells were seeded at ∼80% confluency and treated three days post-infection with Ad-*miR-24*. Cells were stimulated with TGF-β2 (5□ng/mL), IL-1β (0.1□ng/mL), and TNF-α (5□ng/mL) every other day for 6 days. Ad-empty was used as a control.

ARPE-19 cells were transfected with p3xFLAG-MKL1 (Addgene #11978; 1μg) using Lipofectamine™ 3000 (Invitrogen, L3000015) to overexpress MRTF-A. Twenty-four hours later, cells were infected with Ad-*miR-24* or Ad-empty, followed by TGF-β2 (10 ng/mL) treatment 48 h post-infection. For subcellular localization analysis, cells were fixed 24 h after transfection and stained with anti-FLAG antibody (Cell Signaling, 14793, 1:100).

### RNA and Western blot analysis

Total RNA was extracted from ARPE-19 cells using TRIzol™ Reagent (Invitrogen, 15596026). Briefly, cells were lysed with TRIzol™, followed by phase separation with chloroform. The aqueous phase was collected, and RNA was precipitated with isopropanol, washed with 75% ethanol, and dissolved in RNase-free water. RNA concentration and purity were assessed using a NanoDrop™ spectrophotometer. miRNA expression was quantified by real-time reverse transcription PCR (RT-qPCR) using miRCURY LNA SYBR Green PCR Kit (Qiagen, 339345). miRCURY LNA miRNA PCR assays specific for *has-miR-24-1-5p* (Qiagen, YP00204357) and *has-miR-24-3p* (Qiagen, YP00204260) were used. U6 snRNA(V2) (Qiagen, YP02119464) and SNORD44 (Qiagen, YP00203902) served as an internal reference. Relative expression levels were calculated using the comparative Ct (ΔΔCt) method.

Protein was extracted using RIPA lysis buffer (Thermo Scientific, 78430) supplemented with Halt™ Protease and Phosphatase Inhibitor Cocktail (Thermo, 78441). Lysates were centrifuged at 13,000 × g for 10 minutes at 4°C, and the supernatants were collected. Protein concentration was measured using the Pierce™ BCA Protein Assay Kit (Thermo, 23225). Equal amounts of protein were loaded onto 4-20% Mini-PROTEAN TGX Precast Gels (Bio-Rad, 4561094) and transferred to nitrocellulose membranes (Bio-Rad, 1620115). Membranes were blocked with Intercept (PBS) Blocking Buffer (LI-COR, 927-70010) for 1h at room temperature, then incubated with primary antibodies overnight at 4°C. After washing with 1X TBS containing 0.1% Tween-20, membranes were incubated with the appropriate secondary antibodies for one hour at room temperature. Target proteins were detected using the LI-COR Odyssey imaging system. Band densitometry was quantified using the ImageJ software. Antibodies used were as follows: α-Tubulin (Sigma, 05829, 1:5000), α-SMA (Abcam, ab5694, 1:1000), Fibronectin (BD Biosciences, 610078, 1:1000), N-cadherin (BD Biosciences, 610920, 1:1000), CTGF (Cell signaling, 86641, 1:1000), SMAD3 (Abcam, ab28379, 1:1000), SMAD2 (Abcam, 5339, 1:1000), and LIMK2 (Proteintech, 12350-1-AP, 1:500).

To assess actin dynamics, ARPE-19 cells were infected with Ad-*miR-24* or control virus for 3 days and treated with TGF-β2 for 6 days. G- and F-actin fractions were isolated using the G-actin/F-actin In Vivo Assay Kit (Cytoskeleton, BK037). Cells were lysed in pre-warmed LAS2 buffer, clarified, and ultracentrifuged (100,000×g) to separate G-actin (supernatant) and F-actin (pellet). The F-actin pellet was depolymerized according to the manufacturer’s protocol. Equal volumes of G- and F-actin fractions were analyzed by SDS-PAGE and immunoblotting with anti-actin antibody, and band intensities were used to calculate the G/F-actin ratio.

### Laser-induced fibrosis *ex vivo* model

RPE/choroid explants were generated from 8-week-old C57BL/6J mice immediately following laser injury to establish the *ex vivo* fibrosis model. The RPE/Choroid explant was prepared as previously described(37). Eyes were collected from euthanized mice, rinsed twice in 1× PBS, and the connective tissue and anterior segment (cornea and lens) were removed. Each eyecup was cut into four petals, the neural retina was carefully separated, and the remaining RPE/choroid/sclera complex was mounted onto a 6 × 6 mm PVDF membrane with the scleral side facing the membrane. RPE/Choroid explant was cultured in Cytiva HyClone™ Dulbecco’s High Glucose Modified Eagles Medium (HyClone, SH30022LS), 2% FBS, 1% Penicillin-Streptomycin, 1% MEM Non-Essential Amino Acids (Gibco, 11140-50), and 1% GlutaMAX™ Supplement (Gibco, 35050-061). The culture medium was pre-warmed to 37□°C before use.

To evaluate the function of *miR-24* in the laser-induced fibrosis *ex vivo* model, RPE/choroid explants were treated overnight with Ad-*miR-24* or Ad-empty control. Forty-eight hours later, explants were stimulated with TGF-β2 (10 ng/mL) and TNF-α (10 ng/mL) every other day for a total of four days. RPE/choroid flatmount staining was performed as previously described.

### Bulk RNA-seq sample preparation

For RNA sequencing sample preparation, ARPE-19 cells were cultured under long-term differentiation conditions as described. Differentiated ARPE-19 cells were infected with Ad-*miR-24* at a MOI of 2. Following viral transduction, cells were treated with TNT (TGF-β2, 10 ng/mL; TNF-α, 10 ng/mL) for a total of 6 days. Total RNA was extracted using the RNeasy Plus Mini Kit (Qiagen, 74134) according to the manufacturer’s instructions. RNA samples were treated with DNase I to remove residual genomic DNA and subsequently re-purified prior to sequencing. Three independent biological replicates were prepared for each experimental group. RNA-seq was performed by Plasmidsaurus using Illumina sequencing technology with custom analysis and annotation. Additional methodological details are provided in the Supplementary Information.

### MTT assay and gel contraction assay

For the MTT assay, ARPE-19 cells were seeded in 96-well plates and infected with Ad-*miR-24* or control virus. Four days later, MTT reagent (1 mg/mL; Cayman Chemical, 21795) was added and incubated for 4 h at 37 °C. Formazan crystals were dissolved in DMSO, and absorbance was measured at 540 nm to assess cell viability.

Gel contraction was performed as described(76). Type I collagen (Corning, 354236) was titrated with 1 M NaOH to physiological pH (∼7.4), and the minimal volume required for consistent polymerization was used for each batch. ARPE-19 cells transduced with Ad-*miR-24* or Ad-empty and treated with or without TGF-β2 (10 ng/mL) for 48 h were resuspended (1.5 × 10 cells/mL) and mixed with collagen (final concentration 1 mg/mL). The mixture was neutralized, plated (500 µL/well, 24-well plate), and allowed to polymerize for 20 min at room temperature before culture with or without TGF-β2 for 4 days. Gel contraction was quantified by measuring gel area using ImageJ.

### Immunofluorescence for *in vitro* and *in vivo* studies

For *in vitro* studies, cells were seeded on 1% gelatin-coated coverslips, fixed in 4% paraformaldehyde for 30 min at room temperature, and permeabilized with 0.5% Triton X-100 in PBS. After blocking in PBST (0.1% Tween-20) containing 5% horse serum for 1 h, cells were incubated with primary antibodies overnight at 4 °C, followed by fluorophore-conjugated secondary antibodies (1:500) for 1 h at room temperature. Nuclei were counterstained with DAPI, and coverslips were mounted using VECTASHIELD® PLUS Antifade Mounting Medium (Vector Laboratories; H-1900). For ZO-1 staining in differentiated ARPE-19 cells, fixation was performed with cold methanol for 10 min, followed by blocking and antibody incubation as described above. Primary antibody: Fibronectin (BD Biosciences, 610078, 1:100), Collagen III (Abcam, ab7778, 1:200) and ZO-1 antibody (Cell Signaling Technology, #13663,1:100). Secondary antibody: Alexa fluor 488 goat anti-rabbit IgG (Invitrogen, A11001), Alexa Fluor 647 goat-anti-rabbit IgG (Life technologies, A21244), and Alexa Fluor 594 goat-anti-rabbit IgG (Life technologies, A11012).

RPE/Choroid tissue was prepared as previously described(77). Briefly, eyes from 8-week-old mice were enucleated, rinsed in PBS, and the anterior segment (cornea and lens) and neural retina were removed. The remaining RPE/choroid complex was dissected into four radial petals and fixed in 4% PFA overnight at 4 °C. The following day, tissues were washed with PBS and blocked in PBS containing 10% goat serum and 1% Triton X-100 (Sigma, T8787) for 2 h at room temperature. Samples were incubated overnight at 4 °C with primary antibodies diluted in blocking buffer. After six washes with PBS-TX (PBS containing 0.1% Tween-20 and 0.3% Triton X-100), tissues were incubated with fluorophore-conjugated secondary antibodies for 2 h at room temperature. RPE/Choroid flatmount were mounted using VECTASHIELD® PLUS Antifade Mounting Medium (Vector Laboratories; H-1900). Primary antibodies used included periostin (Proteintech, 19899-1-AP; 1:100), α-smooth muscle actin (α-SMA; Abcam, ab5694; 1:200), α-smooth muscle actin (α-SMA; Sigma, C6198) and fibronectin (BD Biosciences, 610077; 1:200), RPE65 (Abcam, ab231782), NG2 (Millipore, AB5320), CD102 (BD Biosciences, 553326), CD45 (BD Biosciences, 553076), Glutamine Synthetase (GS, Abcam, ab73593), Collagen I (Sigma, SAB4500369), Myh11 (Abcam, ab224801), Alexa Fluor™ 594 Conjugate-Isolectin GS-IB4 (Invitrogen, I21413) and rat anti-mouse F4/80 (1:200, a gift from Dr. Philip Thorpe at UT Southwestern Medical Center).

For detection of active collagen accumulation, tissues or cultured cells were fixed in 4% PFA for 1 h, permeabilized with 0.5% Triton X-100, and incubated overnight at 4 °C with fluorescent collagen hybridizing peptide (F-CHP; 3Helix, FLU60) at 10 µM. Prior to use, F-CHP solution was heated to 80 °C for 5 min and rapidly cooled on ice. Samples were incubated with the F-CHP solution overnight at 4 °C. Nuclei were counterstained with DAPI, and samples were mounted using VECTASHIELD® PLUS Antifade Mounting Medium. All images were acquired using a Nikon Eclipse Ti-S/L100 inverted fluorescence microscope or a Nikon A1 laser-scanning confocal microscope and analyzed using Imaris software.

### Image analysis and Statistics

For in vitro immunofluorescence staining, three images were acquired from each biological replicate. Each independent experiment included three biological replicates and was repeated three times (N = 3 independent experiments). Fluorescence intensity was quantified using ImageJ and normalized to cell number prior to statistical analysis. Data from the three biological replicates were averaged within each independent experiment. Images were acquired using identical laser power and exposure settings for each antibody within a given experiment and were subsequently denoised using NIS-Elements (Nikon, Tokyo, Japan). Phalloidin-labeled actin structures were reconstructed and quantified in Imaris (Bitplane/Oxford Instruments, version 10.2.0) using the “Surfaces” and “Filaments” modules, and total filament length was used for statistical analysis.

For RPE/choroid flatmount staining, confocal Z-stack images were acquired to ensure complete coverage of the laser-induced lesion area. Optical sections were collected at 1 μm intervals throughout the entire depth of the tissue. Colocalization between fluorescence channels was quantified using the “Coloc” module in Imaris (Bitplane/Oxford Instruments, version 10.2.0). Briefly, selected fluorescence channels were used to generate a colocalization (Coloc) channel, and the percentage of each original channel overlapping with the Coloc channel was quantified. Three-dimensional (3D) reconstructions were generated from confocal Z-stack images using the “Surfaces” module in Imaris (Bitplane/Oxford Instruments, version 10.2.0). Surfaces were generated for individual fluorescence channels, and overlapping signals were identified by masking intersecting regions to create a new surface representing the colocalized volume. Representative 3D-rendered images were visualized from selected viewing angles and exported using the “Snapshot” module in Imaris. For *in vivo* inhibitor studies, surfaces were generated for each fluorescence channel using the “Surfaces” module in Imaris (Bitplane/Oxford Instruments, version 10.2.0), and the volume of each reconstructed surface was quantified for statistical analysis.

### Statistics

Statistical analyses were performed using GraphPad Prism software. Data are presented as mean ± SEM. Statistical significance between groups was assessed using two-tailed unpaired Student’s t tests, one-way analysis of variance (ANOVA), or two-way ANOVA, as appropriate. For *in vivo* fluorescence quantification, fluorescence intensities were summarized for each experimental group, and extreme values were constrained using an interquartile range (IQR)-based capping approach, in which values exceeding 1.5×IQR from the first or third quartile were replaced with the corresponding quartile thresholds. This procedure reduces the influence of extreme values while preserving the overall data distribution for downstream statistical analyses.

### Study approval

All human sample related study was approved by the Institutional Review Board of University of Texas Southwestern Medical Center (UTSW). Animal studies were conducted following the Association for Research in Vision and Ophthalmology (ARVO) statement for the use of animals in ophthalmic and vision research and were approved by the Institutional Animal Care and Use Committees at Tulane University.

### Data and materials availability

Data and materials availability: The RNA-sequencing data generated in this study have been deposited in the NCBI BioProject database under accession code PRJNA1425742 (https://www.ncbi.nlm.nih.gov/bioproject/PRJNA1425742). The custom scripts used for pathway enrichment analysis (PathN) and downstream RNA-seq data processing are available on GitHub (https://github.com/swanglab/ARPE-19_miR-24_EMT). All data associated with this study are present in the paper or the Supplementary Materials.

## Author contributions

Y.W. performed the experiments and wrote the initial draft of the paper. R.U.V. and Y.H. provided the human AMD and control plasma samples. Y.T., K.G.B., Q.Z., C.D., C.B., E.P., C.A.A., Z.R., J.M. performed experiments, Y.W. and D.B. performed data analysis and bioinformatic analyses. Z.Z. and D.H. provided guidance in human experimental design. S.W. conceptualized and edited the paper.

## Acknowledgements

We would like to thank Drs Alfred S. Lewin from University of Florida, Stryder Meadows, Maria Galazo and Xuebin Qin at Tulane University for providing mouse lines. We also thank Dr. Philip Thorpe at UT Southwestern Medical Center for the rat anti-mouse F4/80 antibody.

## Supplementary materials

MATERIALS AND METHODS

Supplementary Figure 1-12

Supplementary Table 1-4

