## Supplementary Material for "Myofibroblast lineage mapping and inhibiting subretinal fibrosis by targeting SMAD3 and MRTF pathways via microRNA-24 functional study"

**MATERIALS AND METHODS**

**RNA sequencing, data processing, and integrative transcriptomic analysis**

Transcriptome-wide differential gene expression analysis was performed using a 3′ end-counting RNA-seq platform (Plasmidsaurus, USA). Four experimental groups were included: G1, Ad-empty; G2, Ad-miR-24; G3, Ad-empty with TNT; and G4, Ad-miR-24 with TNT, with three biological replicates per group.

**RNA-seq library preparation, sequencing and data processing**

Polyadenylated mRNA was captured from total RNA using an oligo-dT primer incorporating a sample-specific barcode, a unique molecular identifier (UMI), and the Illumina Read 1 sequence. Following second-strand synthesis, double-stranded cDNA was fragmented and tagged by tagmentation to incorporate the Read 2 sequence. Libraries were amplified to add unique dual indices (UDI; i5 and i7) and P5/P7 flow-cell adapter sequences. Sequencing was performed on an Illumina NovaSeq X platform in single-end 100-bp (SE100) mode. Raw sequencing data were demultiplexed using BCL Convert (v4.3.6) and fqtk (v0.3.1). Quality control and read filtering were performed using fastp (v0.24.0), including poly-X tail trimming, 3′ quality-based trimming (minimum Phred score of 15), and removal of reads shorter than 50 bp. Filtered reads were aligned to the human reference genome (GRCh38) using STAR (v2.7.11), with non-canonical splice junctions removed. Resulting BAM files were coordinate sorted using samtools (v1.22.1). PCR and optical duplicates were removed using UMI-based deduplication with UMICollapse (v1.1.0). Mapping quality, strand specificity, and overall alignment performance were assessed using RSeQC (v5.0.4) and Qualimap (v2.3), and a comprehensive quality control report was generated using MultiQC (v1.32). Gene-level expression was quantified using featureCounts (Subread v2.1.1) with strand-specific counting and fractional assignment of multi-mapping reads. Features were defined at the exon and 3′ UTR levels, grouped by gene_id, and annotated with gene biotypes from the reference GTF file.

**Expression normalization and differential expression analysis**

Raw expression values were normalized to Counts Per Million (CPM). To ensure robust analysis, the dataset was filtered to include only "active" transcripts with a mean expression of >1 CPM across all biological replicates. For all downstream calculations, a pseudocount of 0.1 was applied to normalized values to allow for Log_2_ transformation (Log_2_(CPM+0.1) and to handle zero-count observations. The therapeutic effect of miR-24 was evaluated by comparing TNT-treated cells with or without miR-24 overexpression (G3 vs G4). Differentially expressed genes (DEGs) were defined using a stringency threshold of ⏐Log_2_(Fold Change)⏐≥ 0.8 and a minimum expression of 2 CPM in the group with higher abundance. Statistical significance for individual genes was assessed using a two-tailed Student’s t-test (n=3 per group).

**Functional enrichment and pathway analysis**

Functional enrichment was performed on the identified DEGs to determine significantly regulated biological processes. Using the Gene Ontology: Biological Process (GO:BP) database, statistical significance was calculated via a one-sided hypergeometric test (Fisher’s Exact Test) against a background of all expressed genes in the ARPE-19 dataset (N = 15,362). The enrichment score for each pathway was defined as the negative logarithm of the p-value (-Log_10_P). Pathways were subsequently ranked to identify the Top 10 downregulated and Top 10 upregulated signatures.

**Integrated miR-24 target–pathway analysis**

To elucidate the regulatory hierarchy of miR-24, an integrated target-pathway analysis was performed. Potential miR-24 targets were identified through the intersection of significantly downregulated genes (Log_2_FC≤ -0.8) from the RNA-seq dataset with predicted and validated targets obtained from TargetScan 8.0 and miRTarBase. These primary target "hubs" were then mapped to established biological pathways (KEGG and Reactome databases) to identify associated downstream effectors. The resulting regulatory network distinguishes between genes directly silenced by miR-24 and those indirectly downregulated as a consequence of primary target suppression. This integrated approach provides a comprehensive map of miR-24-mediated pathway inhibition, illustrating the cascade from microRNA binding to the collapse of pro-fibrotic signaling programs.

**Transcriptional consistency analysis**

The directional stability of the miR-24 rescue effect was visualized through Z-score normalized expression profiles across all experimental groups. Row-wise Z-scores were calculated based on Log_2_-transformed CPM values to standardize expression variance. "Transcriptional consistency" was confirmed when biological replicates demonstrated a uniform directional shift from G3 toward G1 levels upon treatment with miR-24.

**Supplementary figures:**

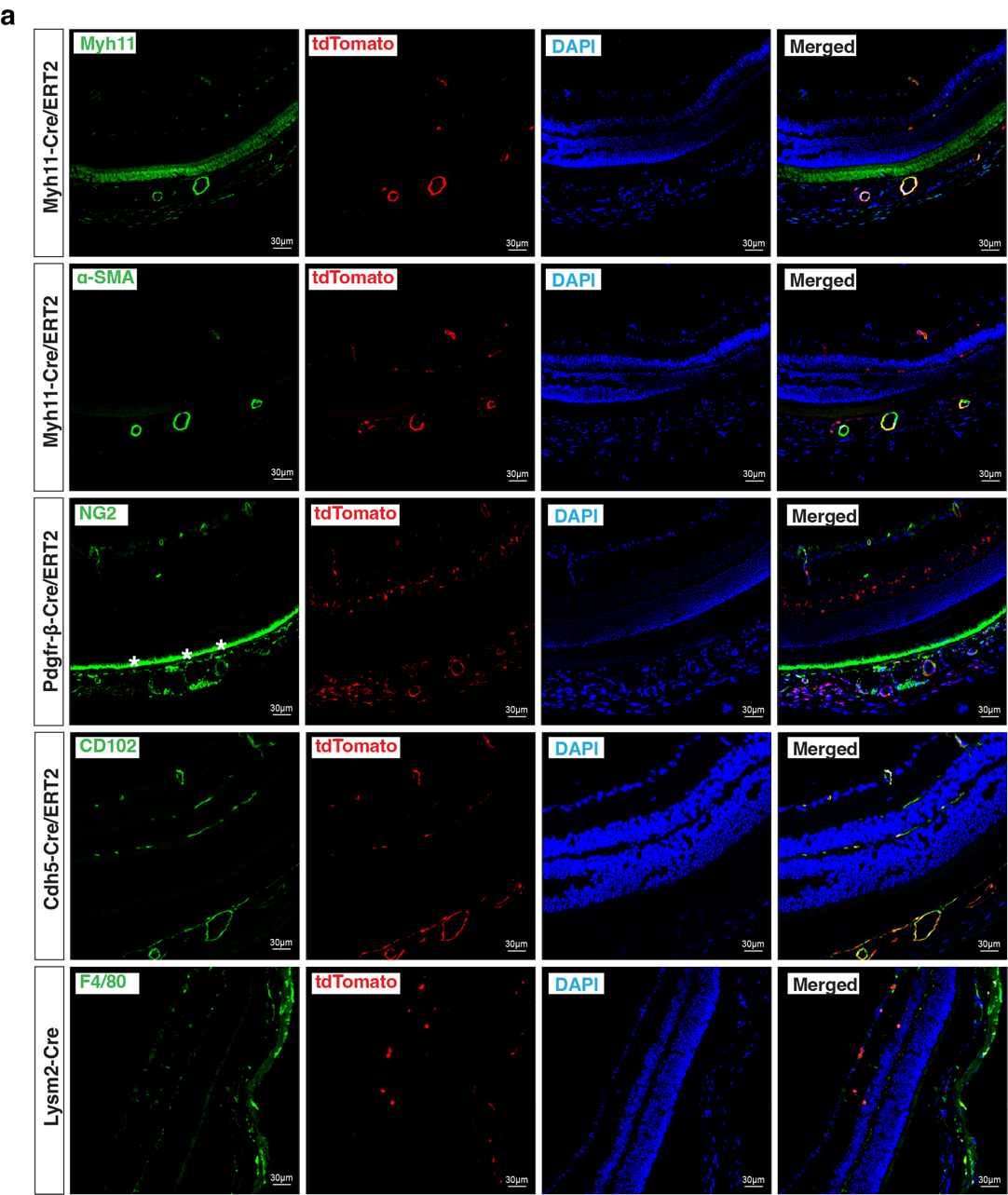

**Supplementary Figure. 1. Validation of lineage specificity in Cre reporter mouse lines under baseline conditions.** Co-staining of tdTomato with cell type-specific markers, including Myh11 and αSMA in the vascular smooth muscles in *Myh11^Cre/ERT2^;R26-tdTomato* mice, NG2 in the pericytes in *Pdgfr-β^Cre/ERT2^;R26-tdTomato* mice, CD102 in the endothelial cells in the *Cdh5^Cre/ERT2^;R26-tdTomato* mice, and F4/80 in the macrophage in the *LysM2^Cre^;R26-tdTomato* mice, confirming the cell type-specificity of the Cre used in the study under baseline conditions without laser injury. n=3.

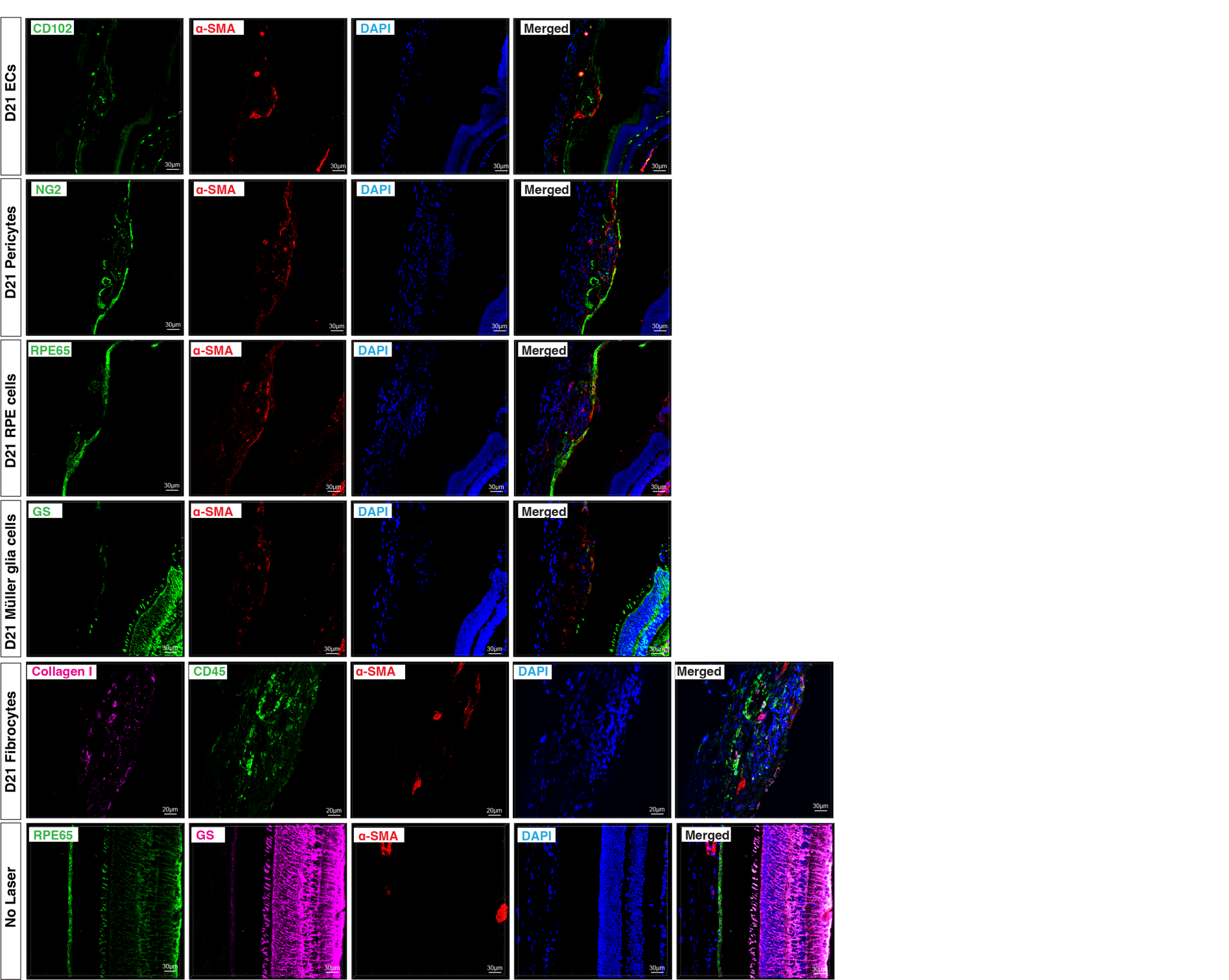

**Supplementary Figure. 2.** Multiple cell types contribute to myofibroblast in laser induced subretinal fibrosis. Co-staining of NG2/α-SMA (pericytes), CD102/α-SMA (ECs), PRE65/α-SMA (RPE cells), GS/α-SMA (muller glia cells), Collagen I/CD45/α-SMA (Fibrocytes) indicating pericytes, ECs and RPE cells can transform into myofibroblast at 21 days after laser injury. n=3.

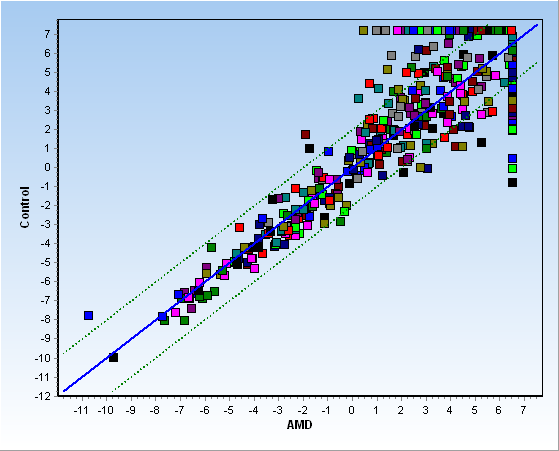

Supplementary Figure. 3. Plasma microRNA profiling in nAMD and control subjects.
Scatter plot showing normalized expression levels of 398 detectable microRNAs in pooled plasma samples from six nAMD patients and six healthy controls analyzed using the miRCURY LNA™ Universal RT microRNA PCR panels. Each dot represents a single microRNA. The solid blue line indicates equal expression between groups, and the dotted green lines indicate a two-fold difference in expression. MicroRNAs outside the dotted lines were considered differentially expressed.

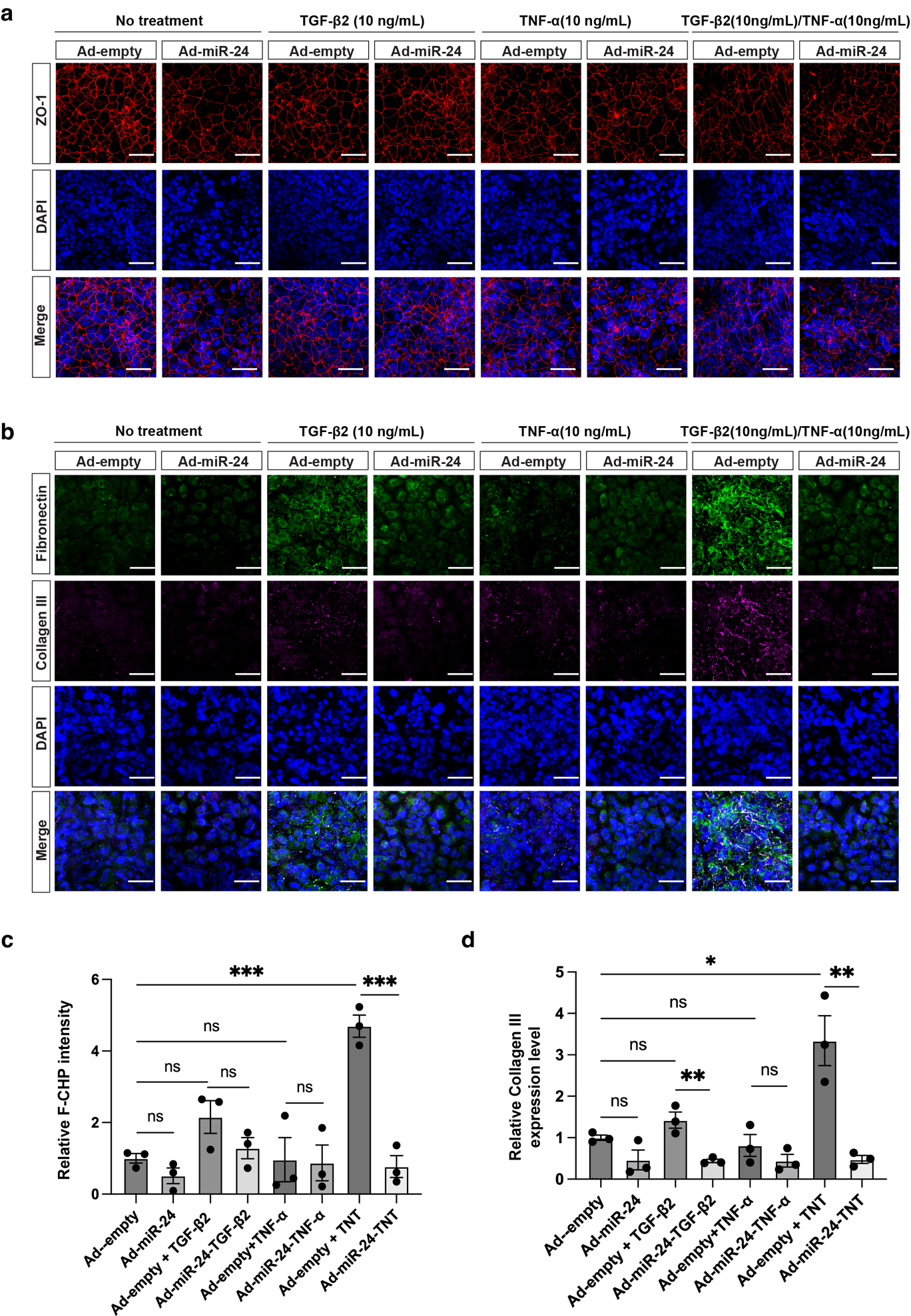

Supplementary Figure. 4. *miR-24* overexpression represses TGF-β2/TNF-α-induced EMT in differentiated ARPE-19 cells. a Representative immunostaining showing that *miR-24* overexpression reduced RPE elongation induced by TGF-β2 (10 ng/mL), TNF-α (10 ng/mL), or their combination in ARPE-19 cells cultured in MEM-Nam medium for 2 weeks. (b-d) Representative immunostaining showing (b) and their quantification (c, d) showing *miR-24* overexpression repressed TGF-β2/TNF-α (TNT)-induced EMT and fibrosis (F-CHP staining and Collagen III) in ARPE-19 cells cultured with MEM-Nam medium for 2 weeks. n=3. *p < 0.05; **p < 0.01; ***p < 0.001; ns, not signiﬁcant.

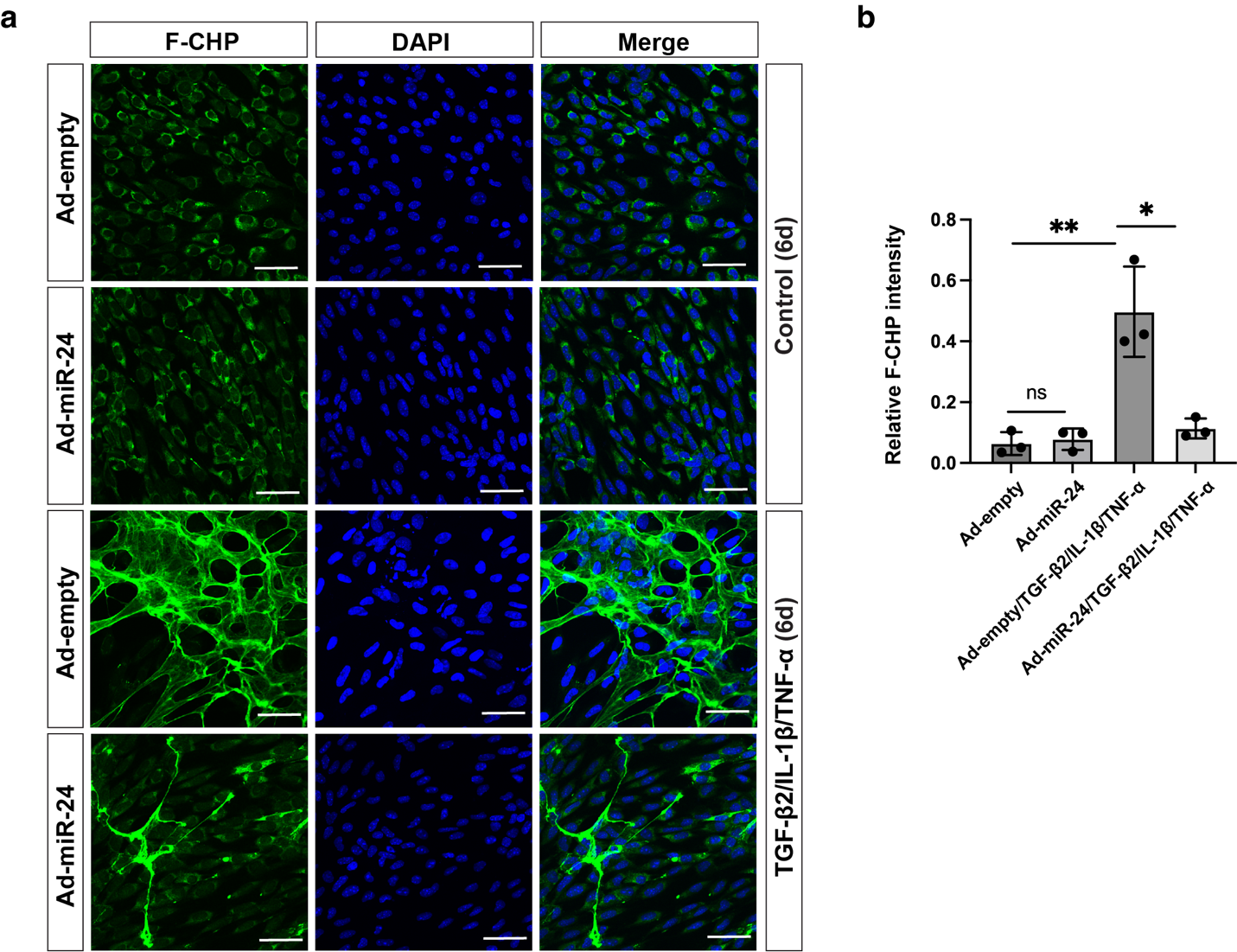

**Supplementary Figure. 5. *miR-24* overexpression represses TGF-β2/TNF-α/IL-1β-induced EndMT in MS1 cells. a** Immunostaining showing that *miR-24* overexpression represses active collagen accumulation, as assessed by F-CHP staining, in MS1 cells treated with TGF-β2/TNF-α/IL-1β every other day for 6 days. **b** Quantification of (a). n=3. *p < 0.05; **p < 0.01; ns, not signiﬁcant.

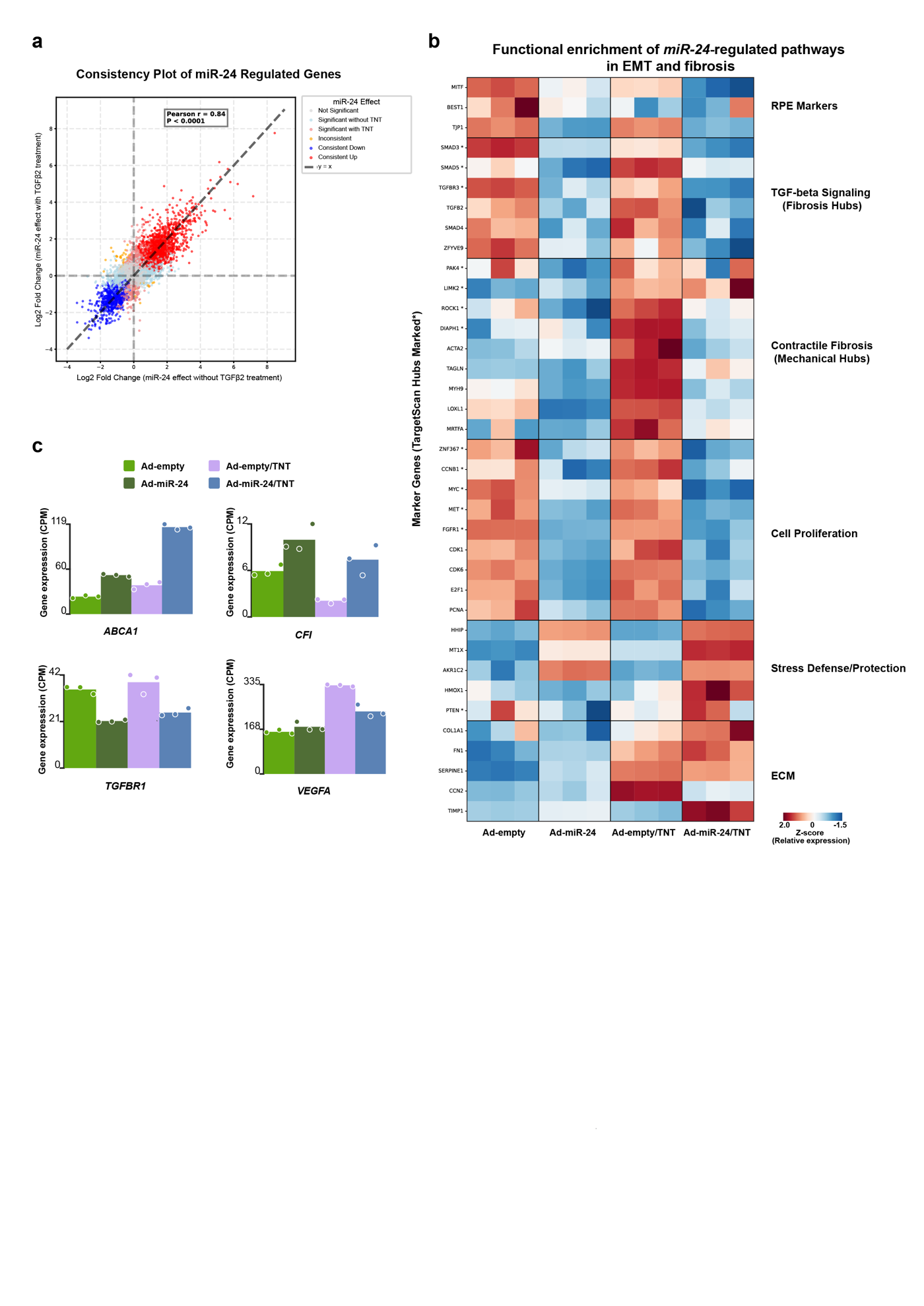

**Supplementary Figure. 6. a** Consistency plot comparing miR-24-induced gene expression changes in ARPE-19 cells in the absence or presence of TNT (TGF-β2 10ng/mL and TNF-α 10 mg/mL). Each dot represents one gene plotted by log₂ fold change with versus without TGF-β2. Genes are color-coded by significance and directionality of miR-24 regulation. Dashed lines indicate zero change and unity; Pearson correlation coefficient and P value are shown. **b** Heatmap of relative expression (Z-score) of selected marker genes and predicted or validated miR-24 targets across Ad-empty, Ad-miR-24, Ad-empty with TNT, and Ad-miR-24 with TNT conditions. Genes are grouped by functional categories, including RPE markers, TGF-β signaling, contractile fibrosis, cell proliferation, stress defense, and extracellular matrix (ECM). Asterisks (*) denote TargetScan-identified hub genes. **c** Bar plots showing normalized expression (CPM) of representative AMD-associated genes, including ABCA1, CFI, TGFBR1, and VEGFA, across experimental conditions. Dots represent biological replicates.

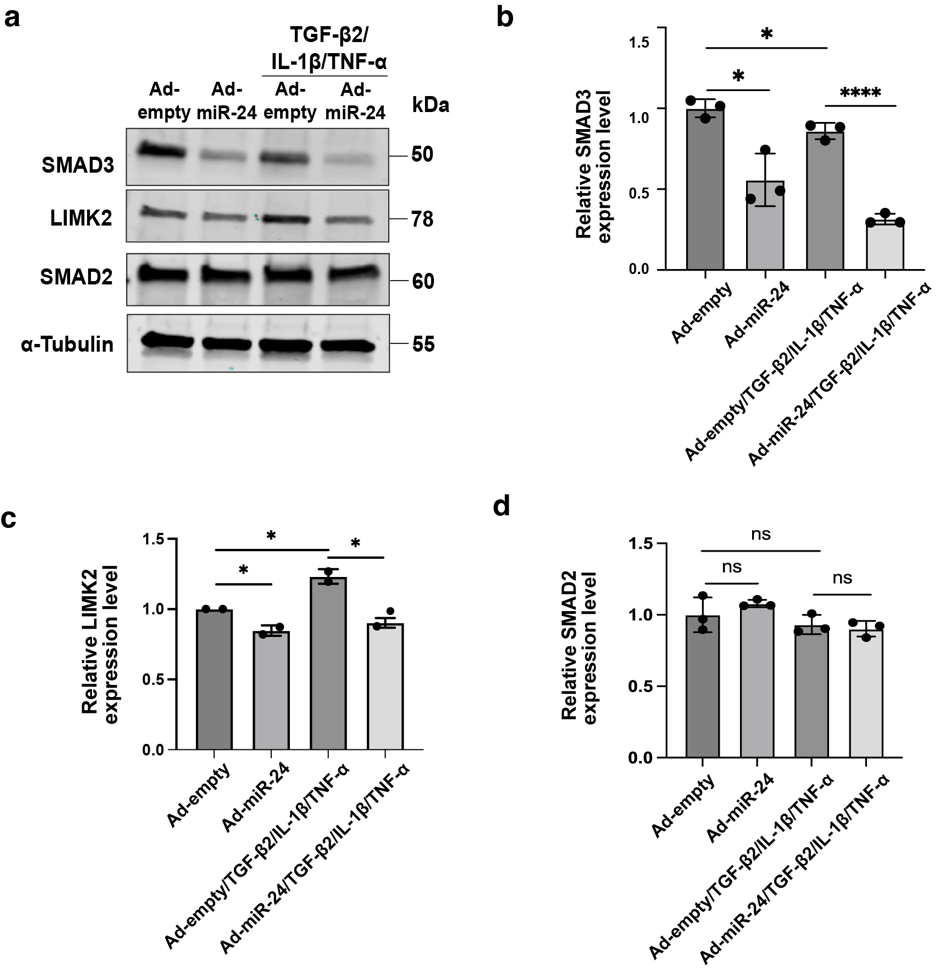

**Supplementary Figure. 7. miR-24 downregulates expression of target genes, SMAD3 and LIMK2, in HUVECs. a** Repression of *miR-24* represses the expression of target protein (SMAD3), and known-target protein (LIMK2), but not SMAD2, by *miR-24* overexpression in HUVECs, with or without TGF-β2/TNF-α/IL-1β treatment for 6 days. **b-d** Quantification of (a). n=3. *p < 0.05, **p < 0.01; ****p < 0.0001. ns, not signiﬁcant.

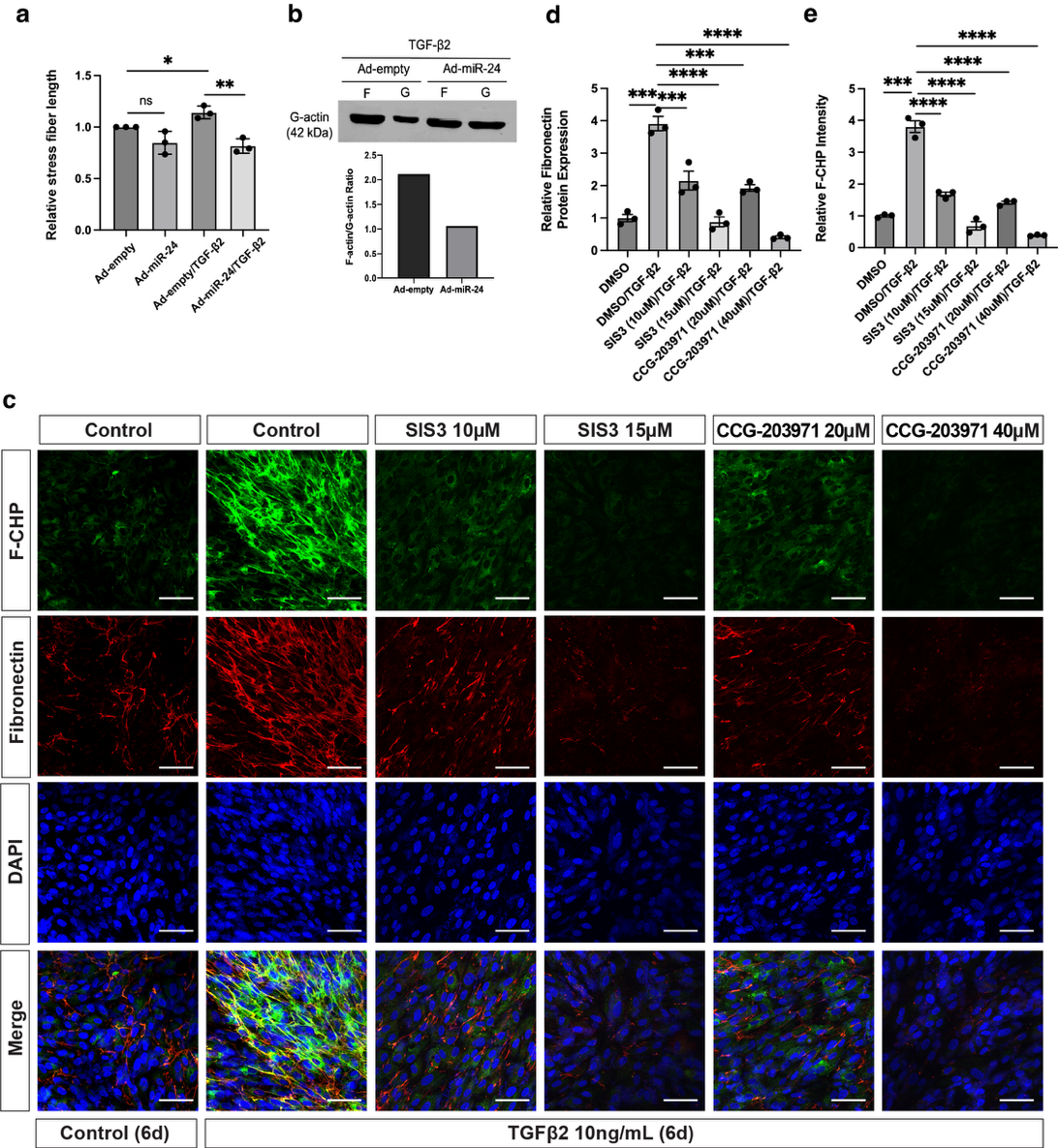

**Supplementary Figure. 8. *miR-24* overexpression suppresses actin cytoskeleton remodeling and EMT through SMAD3 and MRTF-A signaling in ARPE-19 cells *in vitro*.** **a** Quantification of Figure 7**g**, showing that *miR-24* overexpression represses stress fiber formation in ARPE-19 cells treated with TGF-β2 (10 ng/mL) for 6 days. **b** Western blot analysis showing that G-actin formation is increased by *miR-24* overexpression in ARPE-19 cells after TGF-β2 (10 ng/mL) treatment for 6 days. **c** Representative immunostaining showing that higher concentrations of the SMAD3 inhibitor SIS3 (15 μM) and the MRTF-A inhibitor CCG-203971 (40 μM) fully repress EMT and fibrosis in TGF-β2 (10 ng/mL)-treated ARPE-19 cells. **d, e** Quantification of (**c**). n=3. *p < 0.05; **p < 0.01; ***p < 0.001; ****p < 0.0001. ns, not signiﬁcant.

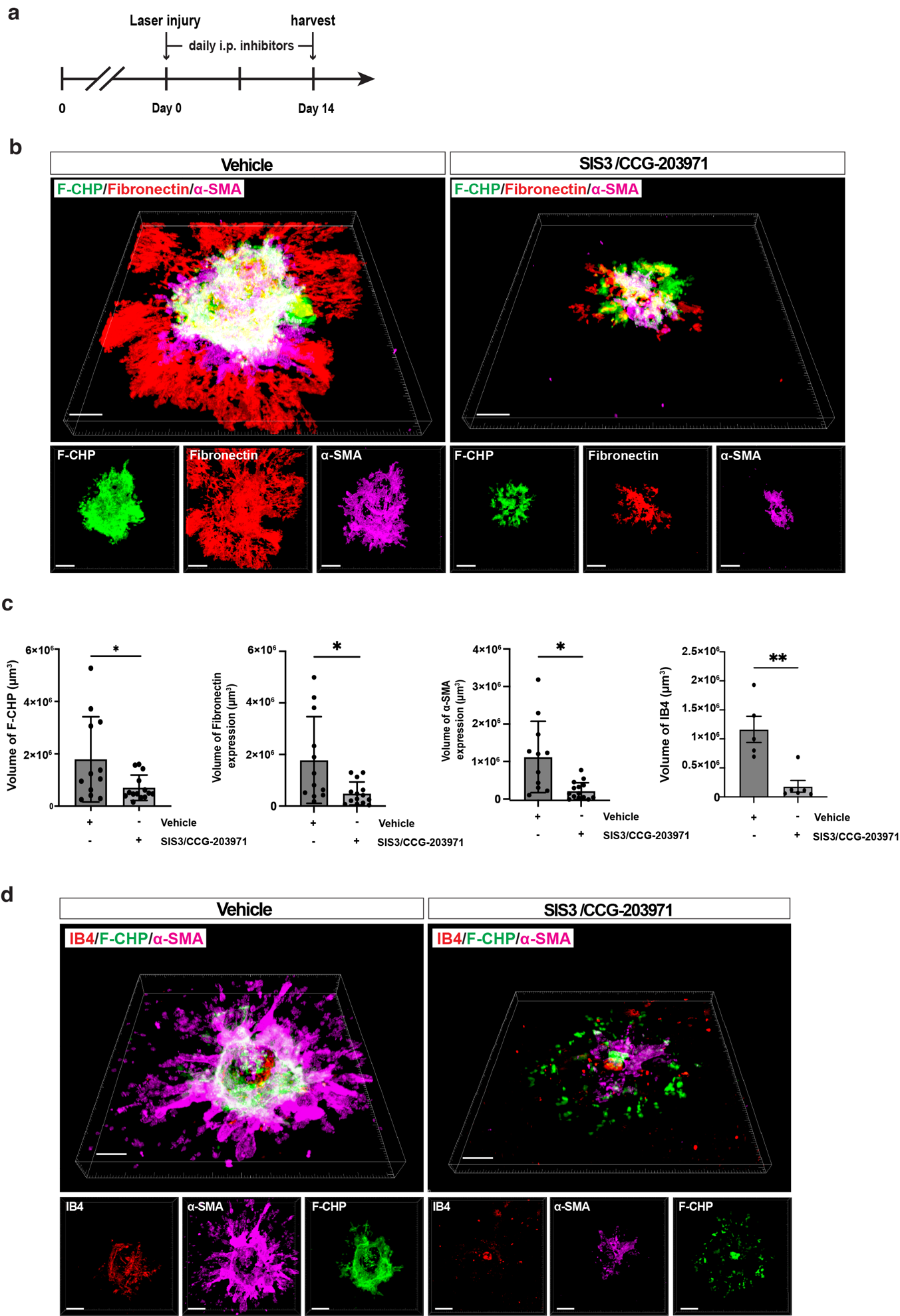

**Supplementary Figure. 9. Early dual inhibition of SMAD3 and MRTF signaling attenuates laser-induced subretinal fibrosis in vivo. a** Experimental timeline for early dual-inhibitors treatment in laser-induced subretinal fibrosis. Combined SIS3/CCG-203971 treatment was initiated immediately after laser injury and administered daily by intraperitoneal injections until day 14. **b** Representative three-dimensional reconstructions of RPE/choroid flatmount stained for F-CHP, Fibronectin, and α-SMA are shown. **c** Quantification of (**b and d**), demonstrating that early dual inhibition significantly reduced F-CHP-positive collagen deposition, α-SMA expression, fibronectin accumulation and CNV formation stained by IB4 compared with vehicle-treated controls. Each dot represents an individual lesion. Data were obtained from 5 mice. *p < 0.05.

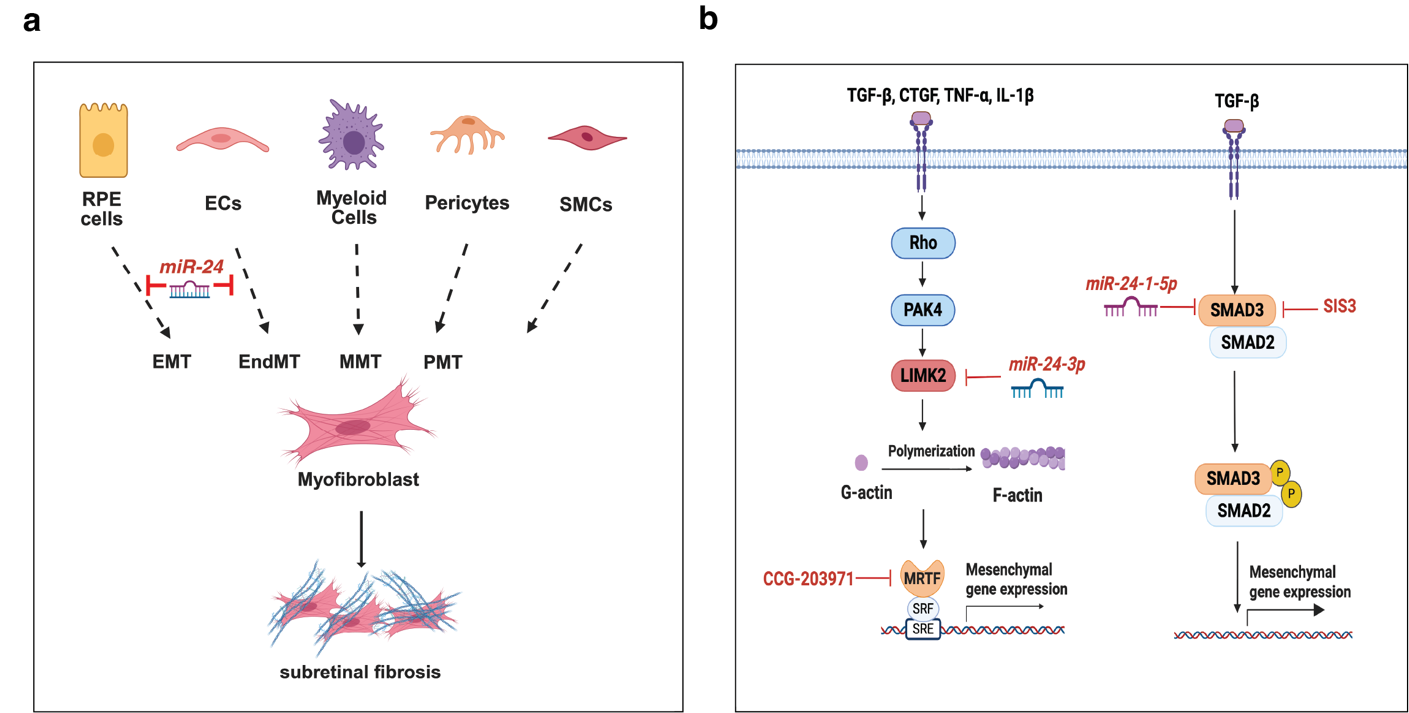

**Supplementary Figure. 10. a working model for *miR-24* in EMT of RPE cells, EndMT of ECs, and subretinal fibrosis. a** RPE cells, ECs, myeloid cells, pericytes, and smooth muscle cells can transform into myofibroblasts, which proliferate and secrete extracellular matrix (ECM), promoting subretinal fibrosis. *miR-24* can repress at least EMT and EndMT to repress subretinal fibrosis. **b** In response to extracellular stimuli (ex., TGF-β, TNF-α, IL-1β and CTGF), Rho signaling activates PAK4, which in turn activates LIMK2 and functions to regulate actin polymerization (F-actin)/ depolymerization (G-actin) dynamics. MRTF-A is a critical co-transcription factor for EMT, EndMT, and fibrosis. As F-actin formation occurs, MRTF-A translocate into the nucleus and regulates mesenchymal gene expression. In the activated TGF-β/SMAD3 signaling pathway, phosphorylated SMAD2/3 and their nuclear transportation will regulate EMT- and EndMT- related gene expression. *miR-24* targets both the TGF-β/SMAD3 signaling pathway and the LIMK2/MRTF signaling pathway downstream of multiple factors by targeting SMAD3 and LIMK2. By regulating both the LIMK2/MRTF and TGF-β/SMAD3 pathways, *miR-24* represses RPE-derived EMT, ECs-derived EndMT, probably macrophage-derived MMT, and therefore fibrosis, which represents an excellent “one drug/multiple targets” model for the treatment of subretinal fibrosis in wet AMD and other fibrotic diseases. Consistently, pharmacological inhibition of SMAD3 activation (SIS3) and MRTF-A signaling (CCG-203971) phenocopies the anti-fibrotic effects of miR-24 overexpression in the subretinal fibrosis model.

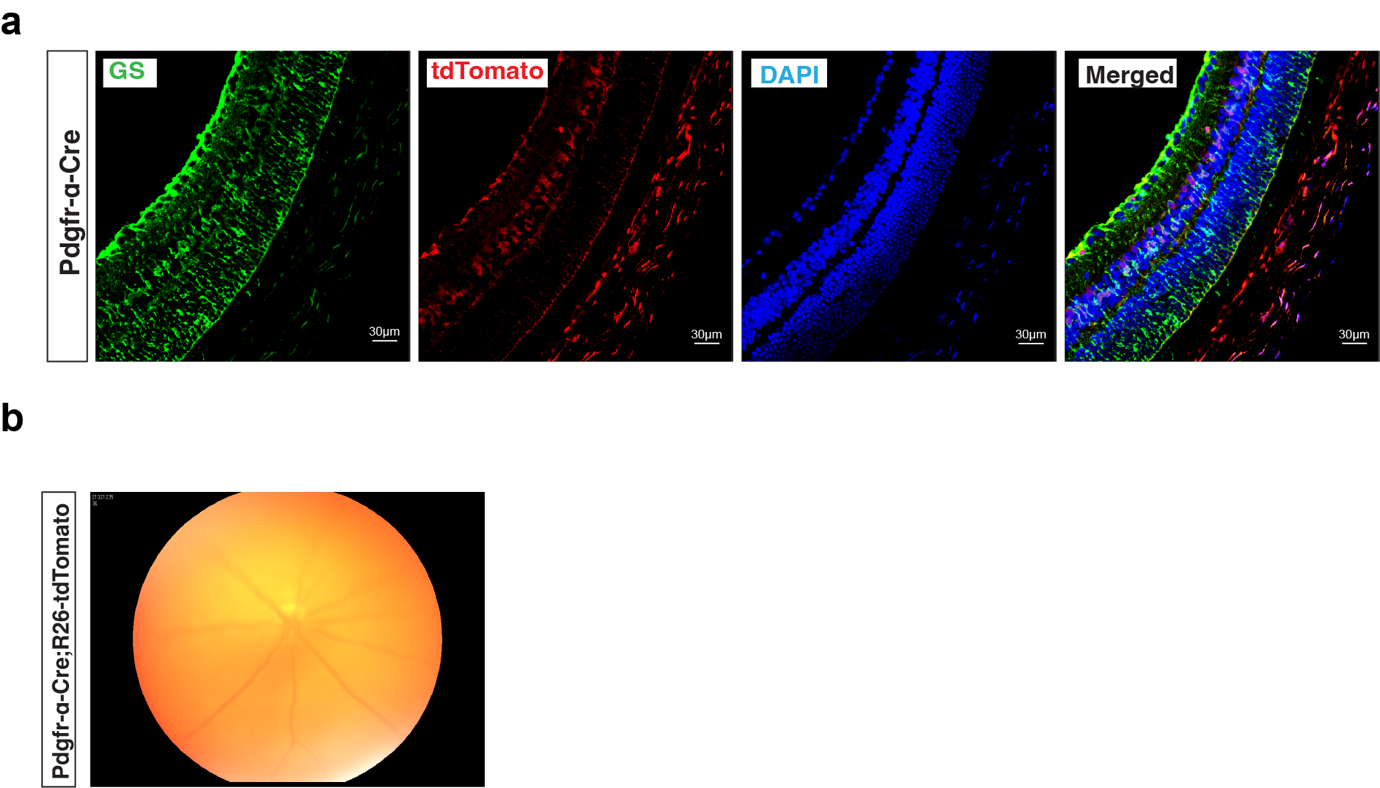

**Supplementary Figure. 11. Validation of Müller glia–specific labeling by *Pdgfrα-Cre* and impaired fundus visibility in *Pdgfrα-Cre;R26-tdTomato* mice. a** Co-localization of GS and tdTomato in retinal sections confirms that ***Pdgfrα-Cre* la**bels Müller glial cells. **b** Representative fundus image of ***Pdgfrα-Cre;R26-tdTomato*** mice showing that widespread tdTomato expression renders the fundus poorly visible.

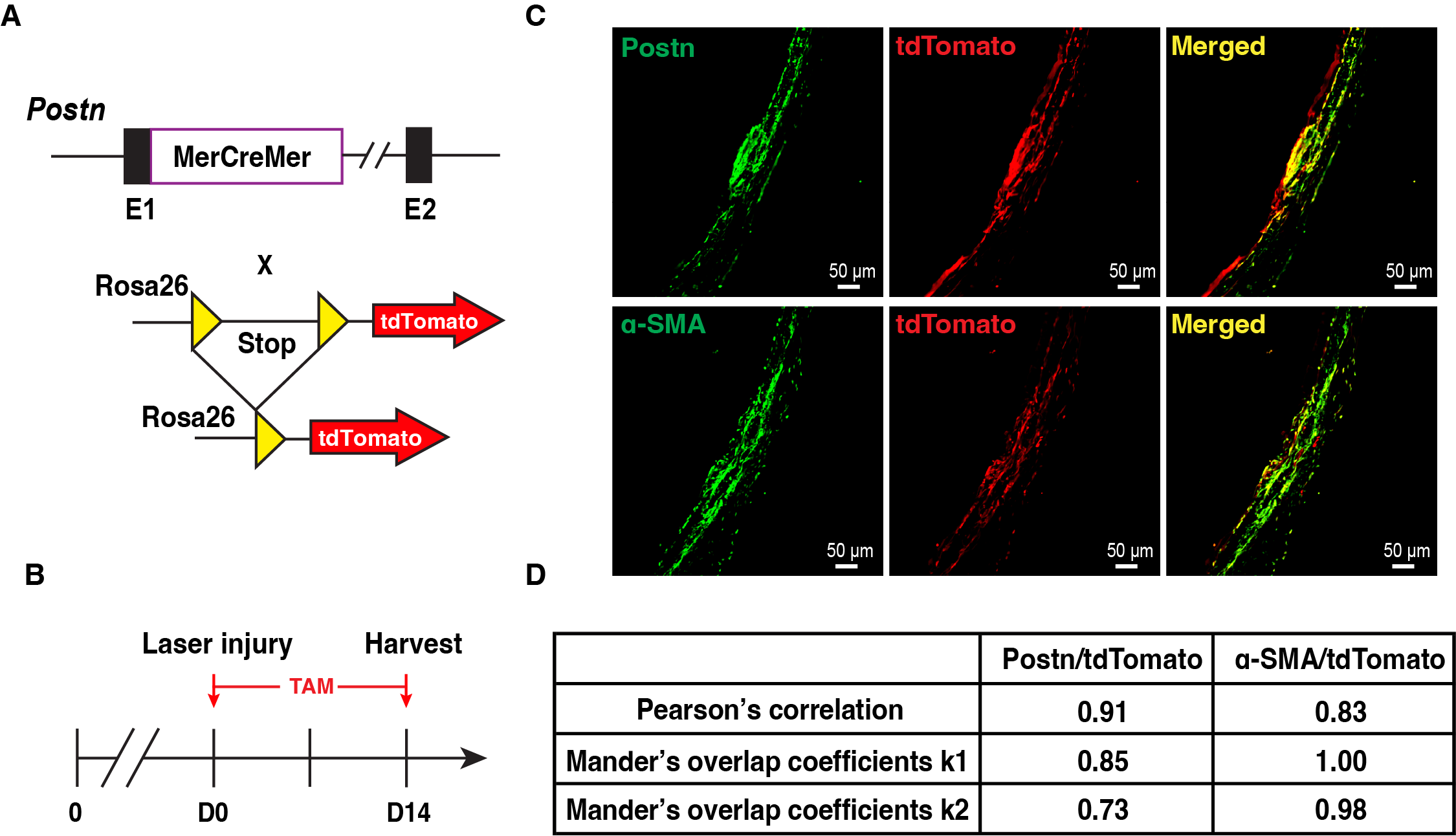

**Supplementary Figure. 12.** Postn labels myofibroblasts after laser injury. **a** Schematics of Postn^+^ cell genetic labeling; **b** Experimental schematic for lineage tracing of myofibroblasts in the laser-induced subretinal fibrosis model. Laser photocoagulation was performed at day 0 (D0), followed by tamoxifen (TAM) administration from D0 to D5 and then every other day by intraperitoneal injection during the indicated period. Eyes were harvested for analysis at day 14 (D14). **c** Representative images of lesion sections showing Postn/tdTomato and α-SMA/tdTomato expression at 14 days after laser injury; Scale bar, 50mm; three laser burns per eye; n=3. **d** Pearson’s correlation and Mander’s overlap coefficients of the signals analyzed by Nikon NIS Elements AR software.

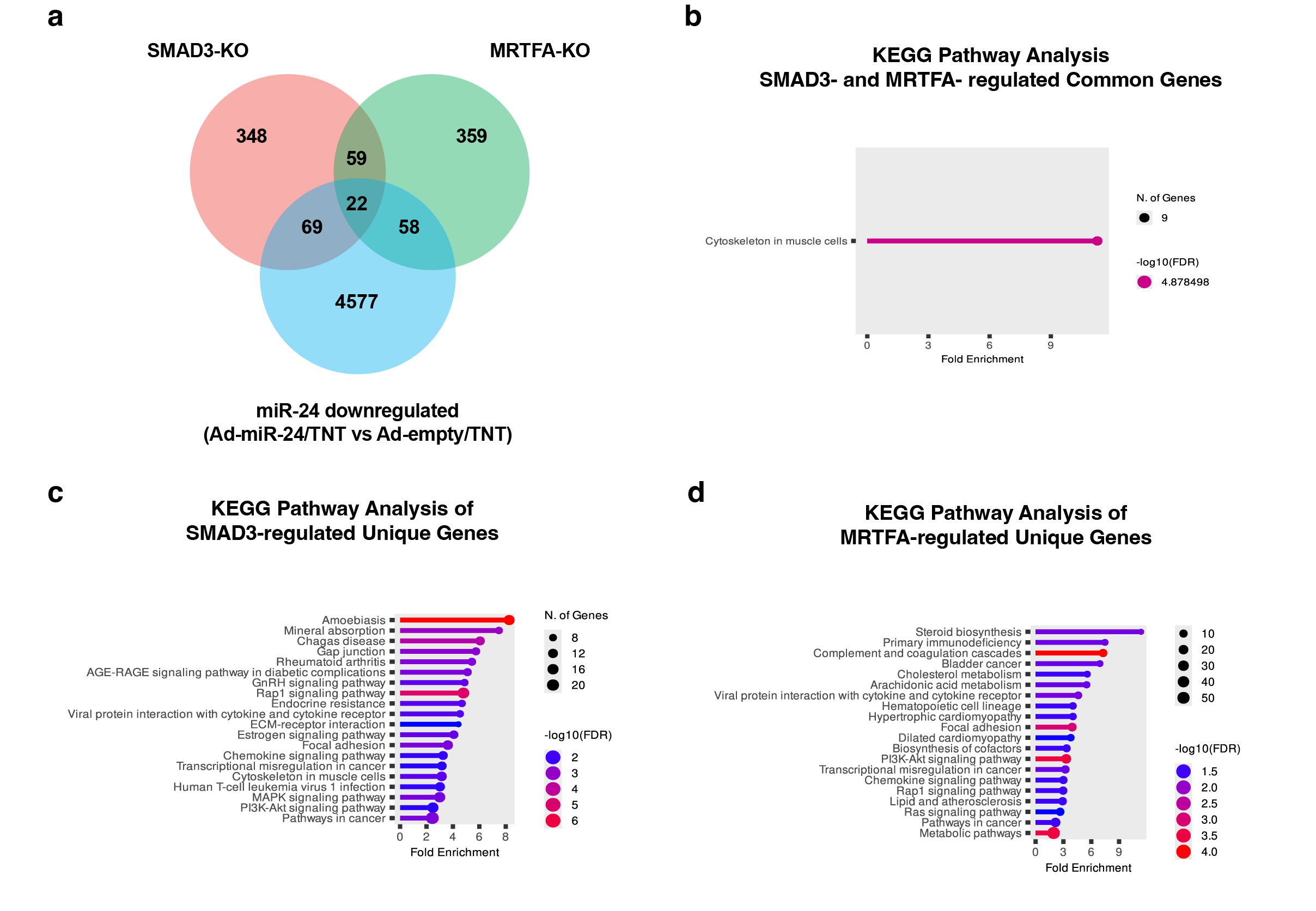

**Supplementary Figure. 13.** **KEGG pathway analysis of SMAD3- and MRTFA-regulated genes.** **a** Venn diagram showing the overlap between SMAD3-regulated and MRTFA-regulated genes. **b** KEGG pathway analysis showing enrichment of pathways among genes commonly regulated by SMAD3 and MRTFA. **c** KEGG pathway analysis of SMAD3-regulated unique genes. **c** KEGG pathway analysis of MRTFA-regulated unique genes. **d** Summary of representative signaling pathways and selected genes uniquely regulated by SMAD3 or MRTFA.

**Supplementary Table 1.** Highly Up and Down Regulated microRNA (|ΔCT_(Control-AMD)_|>2)

| **Up Regulated**  hsa-miR-532-3p  hsa-miR-412  hsa-miR-26a-1*  hsa-miR-147b  hsa-miR-23a*  hsa-miR-409-5p  hsa-miR-544  hsa-miR-501-3p  hsa-miR-520d-3p  hsa-miR-34b*  hsa-miR-642a  hsa-miR-708  hsa-miR-542-5p  hsa-miR-886-3p  hsa-let-7c*  hsa-miR-205  hsa-miR-577  hsa-miR-624*  hsa-miR-23b*  hsa-miR-18a  hsa-miR-885-5p  hsa-miR-876-3p  hsa-miR-605  hsa-miR-920  hsa-miR-505*  hsa-miR-555  hsa-miR-486-3p  hsa-miR-603  hsa-miR-595  hsa-miR-628-5p  hsa-miR-431*  hsa-miR-451  hsa-miR-520h  hsa-miR-181a-2*  hsa-miR-1270  hsa-miR-550a*  hsa-miR-122  hsa-miR-99b*  hsa-miR-629*  hsa-miR-1249  hsa-miR-185*  hsa-miR-129*  hsa-miR-376a*  hsa-miR-658  hsa-miR-1269  hsa-miR-608  hsa-miR-182*  hsa-miR-1913  hsa-miR-1179  hsa-miR-548a-5p  hsa-miR-758  hsa-miR-1183 | **ΔCT**  6.7537  6.2215  5.3589  5.0454  4.9285  4.8812  4.7014  4.5803  4.4359  4.4308  4.40501  4.36032  4.33974  4.16674  4.12506  3.80808  3.76469  3.74396  3.72899  3.66777  3.62008  3.57014  3.52317  3.43604  3.3788  3.28448  3.2581  3.15881  3.14547  3.11736  3.04058  2.98646  2.90773  2.87766  2.85241  2.81566  2.73158  2.69722  2.66159  2.62446  2.60924  2.56373  2.47177  2.32728  2.29822  2.27369  2.25867  2.20511  2.20503  2.11766  2.07962  2.00987 | **Down Regulated**  hsa-miR-187  hsa-miR-301b  hsa-miR-181c  hsa-miR-124  hsa-miR-34a*  hsa-miR-296-5p  hsa-miR-708*  hsa-miR-493  hsa-miR-449a  hsa-miRPlus-D1033  hsa-miR-941  hsa-miR-454  hsa-miR-224*  hsa-miR-760  hsa-miR-200c  hsa-miR-15a*  hsa-miR-30a  hsa-miR-127-5p  hsa-miR-500a  hsa-miR-106b*  hsa-miR-937  hsa-let-7f-1*  hsa-miR-154*  hsa-miR-570  hsa-miR-877*  hsa-miR-643  hsa-miR-146b-5p  hsa-miR-10b  hsa-miR-148b*  hsa-miR-125b  hsa-miR-99b  hsa-miR-940  hsa-miR-664  hsa-miR-29c*  hsa-miR-7-1*  hsa-miR-98  hsa-miR-765  hsa-miR-181c* | **ΔCT**  -7.28707  -6.58579  -6.05091  -4.56316  -4.49444  -4.05252  -3.92602  -3.85482  -3.79389  -3.77421  -3.65106  -3.40043  -3.39847  -3.09284  -3.06356  -3.00732  -2.98286  -2.97964  -2.97502  -2.76811  -2.73425  -2.65233  -2.60209  -2.58074  -2.55754  -2.54048  -2.44293  -2.38879  -2.37829  -2.3285  -2.31842  -2.30371  -2.1928  -2.13648  -2.08344  -2.08063  -2.02583  -2.00612 |
| --- | --- | --- | --- |

**Supplementary Table 2.** Overlap of miR-24 Targets with RNA-seq Downregulated Genes

| Comparison | Ad-miR-24 vs Ad-empty | | Ad-miR-24 (TNT) vs Ad-empty (TNT) | |
| --- | --- | --- | --- | --- |
| miR24- stands | miR-24-1-5p | miR-24-3p | miR-24-1-5p | miR-24-3p |
| Bulk RNA-seq downregulated genes | 3,468 | | 3,187 | |
| Total predicted/validated miR-24 targets | 761 | 1923 | 761 | 1923 |
| Overlapping genes | 208 | 427 | 176 | 379 |
| % of predicted/validated targets in RNA-seq downregulated genes | 27.33% | 22.20% | 23.13% | 19.71% |

**Supplementary Table 2.** **Enrichment of Predicted and Validated miR-24 Targets within the ARPE-19 Transcriptome.** Downregulated genes identified by bulk RNA-seq in ARPE-19 cells following Ad-miR-24 overexpression (± TNT stimulation; TGF-β2 + TNF-α) were intersected with curated lists of predicted and experimentally validated targets of miR-24-3p and miR-24-1-5p. The table summarizes the total number of downregulated genes, total predicted/validated targets, the number of overlapping genes, and the percentage of miR-24 targets represented within the RNA-seq downregulated gene sets under each condition.

**Supplementary Table 3.** Clinical Dysregulation of AMD Risk Genes

| Gene | Full Gene Name | Clinical Status in AMD Patient |
| --- | --- | --- |
| CFI | Complement Factor I | Downregulated |
| ABCA1 | ATP Binding Cassette Subfamily A Member 1 | Downregulated |
| VEGFA | Vascular Endothelial Growth Factor A | Upregulated |
| TGFBR1 | Transforming Growth Factor Beta Receptor 1 | Upregulated |

**Supplementary Table 3.** **Clinical relevance of selected AMD-associated genes.** Summary of representative genes located at established age-related macular degeneration (AMD) risk loci, including their full gene names and reported direction of dysregulation in AMD patient samples. Clinical status is based on published genetic, transcriptomic, and pathological studies of human AMD.

**Supplementary Table 4.** **Distinct signaling pathways uniquely regulated by SMAD3 and MRTF-A**

| SMAD3 Unique Signaling Pathways | Canonical TGF-β signaling | TGFB1, COL1A2, COL6A2, TNC, THBS1, TIMP1 |
| --- | --- | --- |
|  | AGE-RAGE signaling | NOX1, NOX4, TXNIP, CXCL8, CXCL1, CCL18 |
|  | ECM-receptor interaction | Collagens, Laminins |
| MRTFA Unique Signaling Pathways | Ras signaling | IGF1, INSR, PDGFRA, FLT1, VEGFA, LPAR1, PTGER2, SRC, TIAM1, PAK1, FOSL1 |
|  | Complement and coagulation cascades | FGA, FGG, F12, F13A1, TFPI, SERPIND1, CFB, C4BPA, CD59 |
|  | Lipid/atherosclerosis-related pathways | ABCA1, ABCG1, APOC1, APOC2, APOM, CD36, ACTA2, MMP9, ADAMTS5 |

**Supplementary Table 4. Distinct signaling pathways uniquely regulated by SMAD3 and MRTF-A.** Summary of representative pathways and associated genes selectively enriched in SMAD3- or MRTF-A-dependent transcriptional programs. Pathways were identified through differential expression and functional enrichment analyses, highlighting canonical TGF-β and ECM-related signaling downstream of SMAD3, and Ras, complement/coagulation, and lipid/atherosclerosis-associated pathways preferentially linked to MRTF-A.
